# Back to basics: Immunoglobulin germline reference sequences enable investigations and reveal insights into bat-specific immunity

**DOI:** 10.1101/2025.04.10.648212

**Authors:** Ashley B. Reers, Shijun Zhan, Taylor Pursell, Clara Reasoner, Natasha Hodges, Tanya M. Lama, Tony Schountz, Hannah K. Frank

## Abstract

We generated a highly-contiguous, annotated genome of the Jamaican fruit bat, *Artibeus jamaicensis,* including annotated germline immunoglobulin heavy chain (IGH) and light chain (IGL) loci to understand bat B cell receptor repertoires. The bat germline shares many structures and features described in human immunoglobulin loci. However, some features are unique to *A. jamaicensis*, including an expansion of cysteine-rich IGHV genes. To investigate the relationship between the germline IGH locus and expressed B cell receptors (BCRs), we sequenced the BCRs of wild-caught and captive *A. jamaicensis*, finding an enrichment of IGHV3 and IGHV4 genes. Compared to humans, *A. jamaicensis* had shorter CDRH3s and lower levels of somatic hypermutation. Our results demonstrate that while immunoglobulin loci are largely conserved between bats and humans, distinct differences exist in the bat germline, highlighting the need for more detailed genetic characterization of these mammals.

## Introduction

Bats are uniquely able to interact with numerous pathogens of zoonotic importance, seemingly without negative effects. For this reason, bats have been the focus of comparative genomic, transcriptomic, and serological studies that have sought to uncover aspects of the bat immune system that facilitate their unique relationship with pathogens ^1–9^. While these studies have made clear that bats mount a humoral response to infection involving the production of antibodies by B cells, the specifics of these responses remain poorly understood. Dramatic variability in antibody responses among individuals and species broadly, as well as in response to various pathogens, highlights how little we know about the bat humoral immune response ^7,9–11^. Furthermore, a lack of basic knowledge of baseline B cell characteristics in bats has hampered more targeted studies of humoral immunity.

B cell receptors (BCRs) (and their associated antibodies), in mammals, are encoded in the germline by the immunoglobulin heavy chain (IGH) locus and either the immunoglobulin light chain kappa (IGK) or light chain lambda (IGL) locus ^12–14^. These unique loci contain numerous variable (V), diversity (D), and joining (J) genes that are recombined in a structured fashion to produce a single V-D-J (IGH) or V-J (IGK; IGL) segment ^13,14^. These segments comprise the variable region of the BCR that is responsible for antigen recognition. When expressed, the variable region is associated with a constant, Fc, region (encoded by C genes) that can interact with Fc receptors expressed by other immune cell populations to facilitate pathogen clearance ^12,14^. A significant portion of the diversity of the expressed BCR repertoire is encoded by these germline loci ^14^. Thus, determining the composition of the germline IGH, IGK, and IGL loci provides valuable information about the structure of the B cell response. To date, researchers have annotated the germline IGH loci of only a few bat species (*Rousettus aegyptiacus*, *Eptesicus fuscus*, *Rhinolophus ferrumequinum*, *Phyllostomus discolor*, *Pipistrellus pipistrellus*) and investigated the expressed BCR repertoires of even fewer species *(Myotis lucifugus*, *Pteropus alecto*) (see ^2,5,15–17^). Only one study has investigated the bat germline immunoglobulin light chain loci, in *E. fuscus* ^16^. These studies concluded that B cells have undergone less somatic hypermutation (SHM) than in humans or mice, suggesting bats may rely on germline IGH repertoire diversity rather than SHM when generating antibodies in response to infection ^5,18^.

Here, we characterize the germline immunoglobulin loci of *Artibeus jamaicensis* (Jamaican fruit bat), a species frequently used in immunological studies ^19–21^, to better understand the genetic basis of B cell responses in bats and provide a critical resource for the continued study of bat humoral immunity. To this end, we generated a highly-contiguous *A. jamaicensis* genome assembly that captured the difficult-to-sequence IGH locus on one contig and the IGL locus on two contigs (no IGK locus was identified), allowing for detailed annotation of immunoglobulin genes and regulatory regions. With this germline annotation, we demonstrate the importance of bat-specific or species-specific immunoglobulin references for the accurate characterization of bat BCR-repertoires. Additionally, we applied our IGH reference to assess baseline characteristics of the expressed BCR repertoire in *A. jamaicensis*, revealing lower SHM levels and shorter CDRH3 sequences than those observed in humans. Overall, this study highlights unusual features of the *A. jamaicensis* immunoglobulin loci and expressed BCR repertoire that may influence the adaptive immune response in these bats while demonstrating the importance of genetically characterizing these mammals.

## Results

### Highly-contiguous *Artibeus jamaicensis* genome assembly

The repetitive nature coupled with the length of the IGH locus has made this region difficult to sequence and assemble. In a recent, reference-quality *A. jamaicensis* genome assembly (CSHL_Jam; GCF_021234435.1) this locus is spread over multiple contigs (**Figure S1A**), making it difficult to accurately annotate ^22^. We generated a highly-contiguous genome assembly through PacBio HiFi sequencing technology which generates long reads (10-25 Kb) with base-level resolution (>99.5%) on par with short-read technologies ^23^. We sequenced kidney fibroblast cells from a male *A. jamaicensis* from a captive colony and generated a partially phased assembly with Hifiasm ^24,25^. The resulting assembly captured the genome in 202 contigs with a contig N50 value of 100 Mbp (**Figure 1A**). Based on the BUSCO assessment of our assembly, the gene set is highly complete at 95.3% (94.1% single copy, 1.2% duplicated, 0.8% fragmented, 3.9% missing). This assembly is of similar quality to published genomes for other bat species (**Figure 1A-B**). We also generated a whole genome annotation using TOGA (Tool to infer Orthologs from Genome Alignments) ^26^. TOGA identified 19,305 genes (92.7% intact, 3.6% lost, 3.6% partial or uncertain loss). Protein BUSCO analysis confirmed our whole genome annotation was also highly complete, with 99.7% of BUSCOs complete (43.6% single copy, 56.1% duplicated) and 0.3% missing.

**Figure 1.**
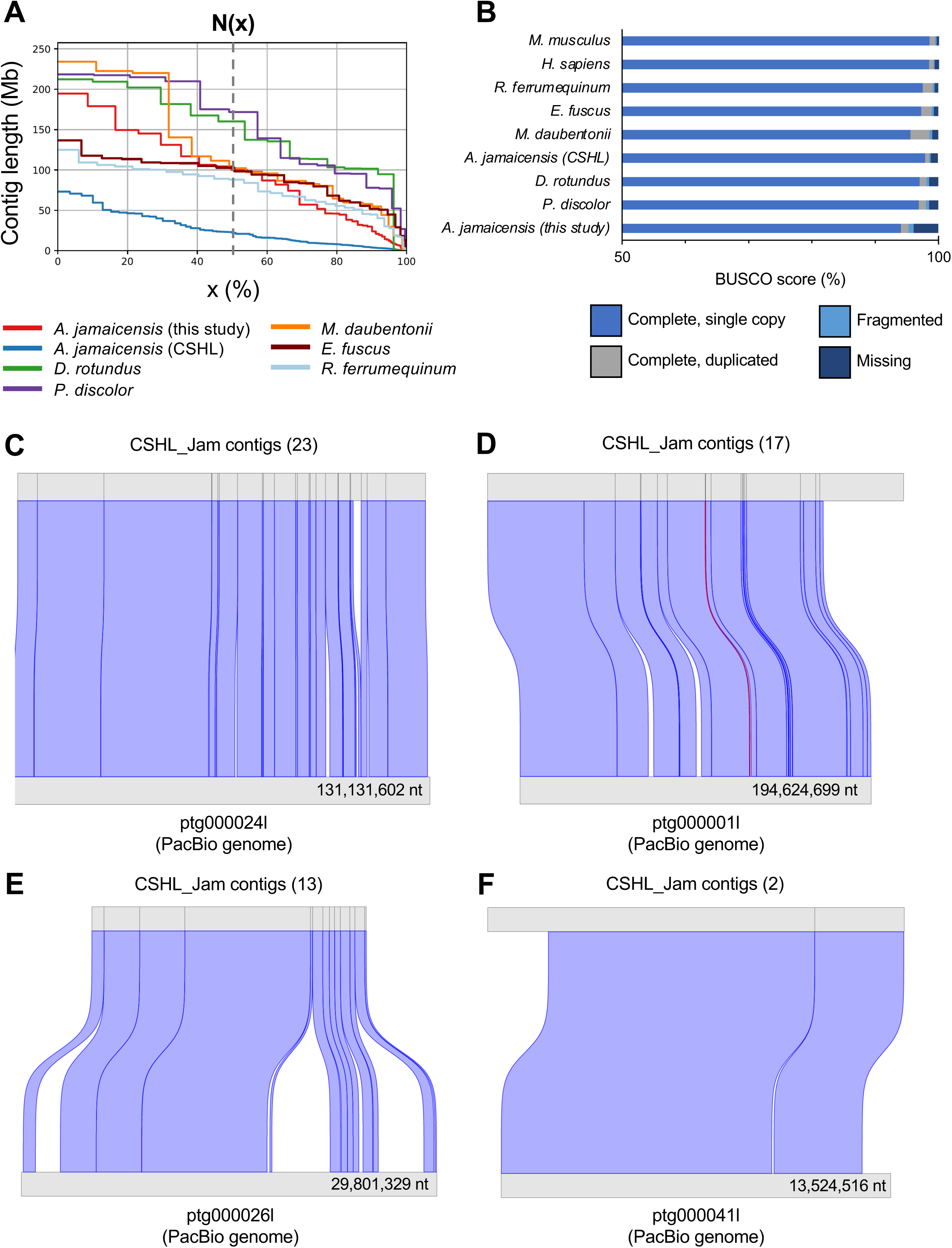
Genome assembly metrics. **(A)** Graph depicting N(x) values for the indicated bat genome assemblies. Dashed gray line indicates the N50 metric. Genome metrics calculated using Quast. **(B)** Assembly BUSCO scores for human, mouse, *Artibeus jamaicensis*, and all chromosome-level bat assemblies with reported BUSCO information (obtained from NCBI). **(C-F)** Synteny plots showing conserved regions between contigs from the CSHL_Jam assembly ^22^ (top) and our PacBio genome assembly (bottom) for **(C-D)** two long contigs from our PacBio assembly, as well as the contigs from our assembly containing the **(E)** IGH locus and **(F)** IGL locus. Alignments were generated with minimap2 and plotted with the R package asynt. Only alignments with a sequence identity >70% and an overlap >100,000 nt were plotted. Blue connections indicate alignments overlapping in the forward direction while red connections indicate inverted alignments. Numbers in parentheses indicate the number of CSHL_Jam contigs aligned to each contig from our assembly.

To understand how our assembly compared to the recently published CSHL_Jam assembly, we assessed the synteny between these assemblies. The two assemblies shared significant sequence identity (**Figure 1C-F**) over the contigs in our PacBio genome containing the IGH and IGL loci. The agreement between these two independent assemblies suggests we have accurately captured the *A. jamaicensis* genome.

The BCR locus of *Artibeus jamaicensis* is similar to other mammals, with species- and bat-specific features.

#### IGHV genes

Using IgDetective, a computational tool that identifies immunoglobulin genes based on the presence of regulatory sequences ^27^, we identified the contig ptg000026l in our assembly as containing the IGH locus (**Figure S1B**). We next identified IGHV, IGHD, IGHJ, and IGHC genes by aligning all reference sequences for these genes available from IMGT and the predicted *de novo* IGHV genes identified by IgDetective to contig ptg000026l ^28^. Recombination signal sequences (RSSs) were annotated for each IGHV, IGHD, and IGHJ gene. For IGHV genes, leader sequences were annotated and used to determine functionality which was assigned based on IMGT definitions ^29^. Using this approach, we identified 196 total IGHV genes (57 functional, 5 ORFs, 134 pseudo) (**Figure 2A, Table S1**). IGHV genes belonged to all three clans and included representatives of families IGHV1 (13 functional, 2 ORF, 33 pseudo), IGHV2 (1 functional, 2 pseudo), IGHV3 (19 functional, 1 ORF, 57 pseudo), IGHV4 (9 functional, 2 ORF, 31 pseudo), IGHV6 (15 functional, 16 pseudo), and IGHV7 (4 pseudo); however, functional IGHV genes were only identified for families IGHV1-4 and IGHV6 (**Figure 2C**). Functional IGHV genes found in *A. jamaicensis* clustered with functional human IGHV genes from the same families (**Figure 2D**).

**Figure 2.**
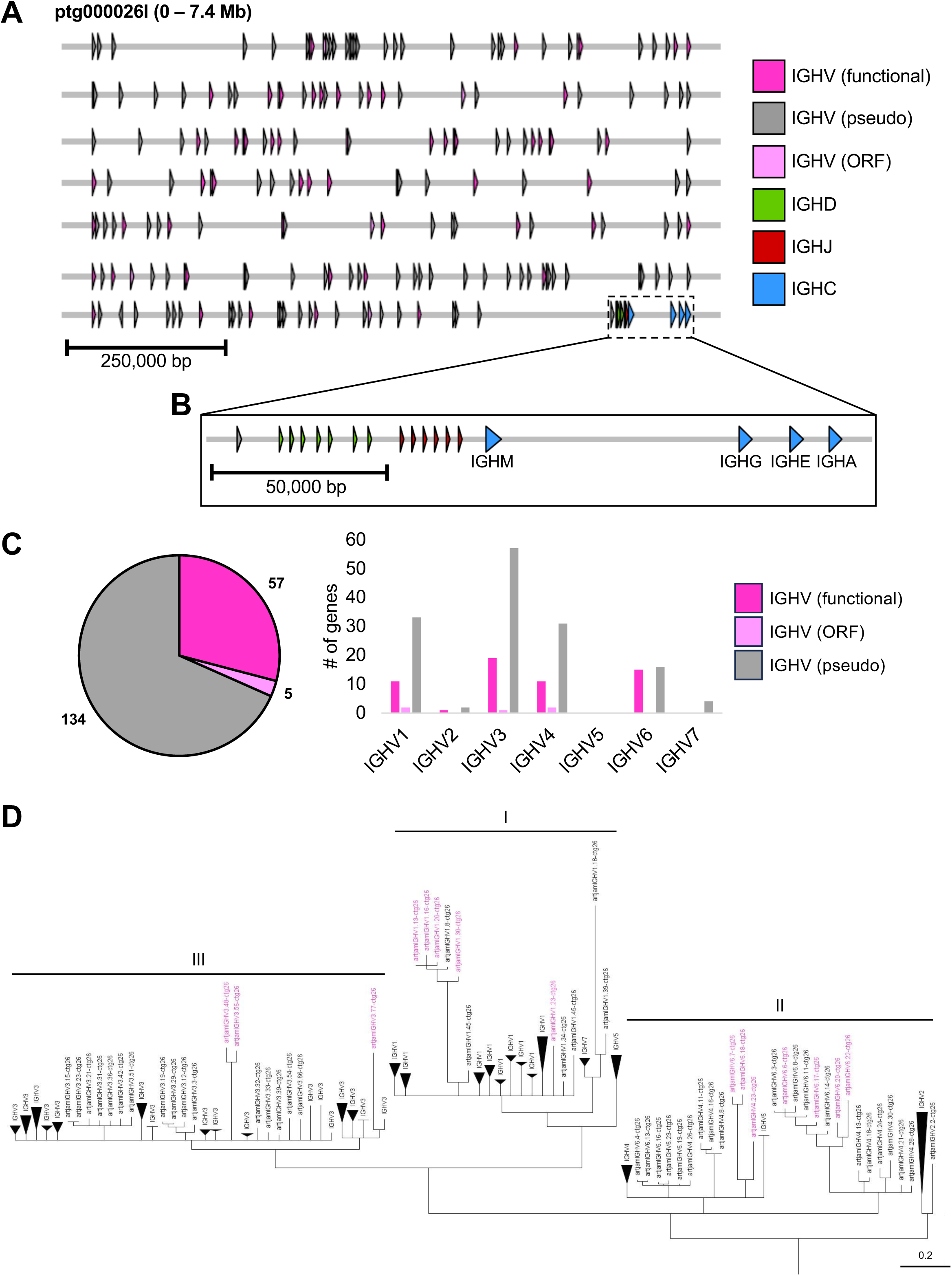
Gene organization in the *A. jamaicensis* IGH locus. **(A)** Schematic overview of the *A. jamaicensis* IGH locus on contig ptg000026l. Arrows indicate directionality of the annotated genes. **(B)** Inset depicts organization of IGHD, IGHJ, and IGHC genes. **(C)** Number of total functional, open reading frame (ORF), and pseudo IGHV genes annotated (pie chart) and within each IGHV family (bar chart). **(D)** Cladogram of all functional *A. jamaicensis* IGHV genes. A single pseudogene (artjamIGHV4.23-ctg26p) was included as it contained extra cysteine residues. Functional human IGHV genes (without artjam prefix) are included for comparison. IGHV genes with 3 or more cysteine residues are shown in pink.

#### IGHD and IGHJ genes

We identified 7 IGHD and 6 IGHJ genes (**Figure 2B, Table S1**). All IGHJ genes appeared functional and contained the WGXG motif conserved in IGHJ genes across species ^28^. IGHD and IGHJ genes were ordered in the locus with all the IGHD genes preceding the IGHJ genes as seen in the human and mouse IGH loci (**Figure 2B**). This organization of IGHD and IGHJ genes has also been reported in *E. fuscus*, *R. ferrumequinum, P. discolor,* and *P. pipistrellus* ^15,16^. Interestingly, in *R. aegyptiacus*, IGHD and IGHJ genes are intermixed in a IGHD-IGHJ-IGHD-IGHJ fashion ^2^. *Rousettus aegyptiacus* and *R. ferrumequinum* belong to the suborder Yinpterochiroptera, while the other bat species belong to the suborder Yangochiroptera. In other aspects of immunity, bat families or species differ; for example, some pteropodid bats have an expansion of interferon genes that has not been seen in yangochiropteran families ^3^, while *Pteropus alecto*, another pteropodid has a contracted type I interferon locus ^30^. The difference in IGHD-IGHJ organization between *R. aegyptiacus* and these other species could represent a species-specific or family level difference. The increasing availability of chromosome-level genome assemblies and IGH locus annotations for more species from both suborders will allow for insights into whether there is additional diversity in IGHD-IGHJ organization.

#### IGHC genes

Within the constant region of the *A. jamaicensis* IGH locus, we identified representatives of all major IGHC isotypes, excluding IGHD (**Figure 2B, Table S1**), consistent with *A. jamaicensis* transcriptomic data and reports from other bat species ^2,15^. To gain insight into the functionality of the annotated IGHC genes, we compared N-glycosylation sites and Fc receptor binding motifs found in humans to those in *A. jamaicensis* (**Figure S2**). *A. jamaicensis* IGHM contained all N-glycosylation sites observed in human IGHM, as well as an additional site in the CH2 region (**Figure S2A**). In IGHG, interaction with FcγR1 is mediated by several critical contacts: the N-glycosylation site at Asn^297^, the lower hinge region (Leu-Leu-Gly-Gly), the CH2 BC loop Asp^265^, and the CH2 FG loop (Ala-Leu-Pro-Ala-Pro) ^31–34^. The *A. jamaicensis* Asn^297^ N-glycosylation site and Asp^265^ were conserved with humans and *E. fuscus*; however, the other binding motifs shared only partial sequence identity with humans (**Figure S2B**). *E. fuscus* also differed from human sequences in these motifs, and this variability in bat IGHG sequences could reflect broader differences in Fc receptors between bats and humans that change the interaction between IGHG and FcγR1. In *A. jamaicensis* IGHE and IGHA sequences, we observed variability in both the Fc receptor binding motifs and the N-glycosylation sites that likely impact receptor binding and effector functions (**Figure S2C-D**). In both IGHE and IGHA, *A. jamaicensis* appeared to have lost most of the N-glycosylation sites present in humans but gained several novel N-glycosylation sites, some of which were shared with *E. fuscus* (**Figure S2C-D**) and appeared to be a feature of these isotypes in bats. Determining how this impacts Fc effector function will require further characterization of bat Fc receptors and their interactions with the full Fc.

To further investigate the functionality of the annotated *A. jamaicensis* constant genes, we searched the constant region for the canonical AID hotspot motif 5’-AGCT-3’ ^35,36^. Increased density of this motif upstream of constant genes facilitates class switch recombination ^35,37^. In keeping with the annotated IGHC genes being functional, all four had dramatic increases in switch motif density directly upstream of the constant genes (**Figure S3A**). To verify expression of the annotated constant genes, we used BLAST to search published transcriptomic data from an uninfected *A. jamaicensis* spleen sample ^21^ for sequences with similarity to the annotated genes. For all IGHC genes, BLAST identified transcripts that aligned with high identity and covered nearly the entire coding region (**Figure S3B**). Fewer transcripts were identified with similarity to IGHE coding regions, consistent with lower IGHE expression in the spleen compared to other antibody isotypes (**Figure S3C**).

#### IGL locus

With limited exceptions (i.e. camelids ^38,39^), a complete, functional BCR requires sequences from both the IGH locus and either the IGK or IGL locus. IgDetective identified IGL genes on contigs ptg000034l and ptg000041l, but did not identify any IGK genes (**Figure S1B)**. We suspect this is a true loss of IGK function; within bats IGK transcripts have only been documented in yinpterochiropteran species and the IGK locus has been lost in *E. fuscus* ^4,16^.

Using the same approach as the IGH locus annotation, we identified 151 IGLV genes (127 functional, 2 ORFs, 22 pseudo) (**Figure 3A-B, Table S1**). IGLV genes belonged to all five clans and included representatives of families IGLV1 (27 functional, 1 pseudo), IGLV2 (3 functional), IGLV3 (10 functional, 1 ORF, 3 pseudo), IGLV4 (5 functional, 1 ORF, 1 pseudo), IGLV5 (11 functional, 1 pseudo), IGLV6 (3 functional), IGLV7 (24 functional, 4 pseudo), IGLV8 (9 functional), IGLV9 (1 functional), IGLV10 (1 functional), and IGLV11 (33 functional, 12 pseudo) (**Figure 3B**). Functional IGLV genes clustered with human IGLV genes from the same families, and genes from both contigs clustered together (**Figure 3C**). Since the IGL locus was split over 2 contigs, we cannot rule out the existence of additional IGLV genes not captured by our genome assembly.

**Figure 3.**
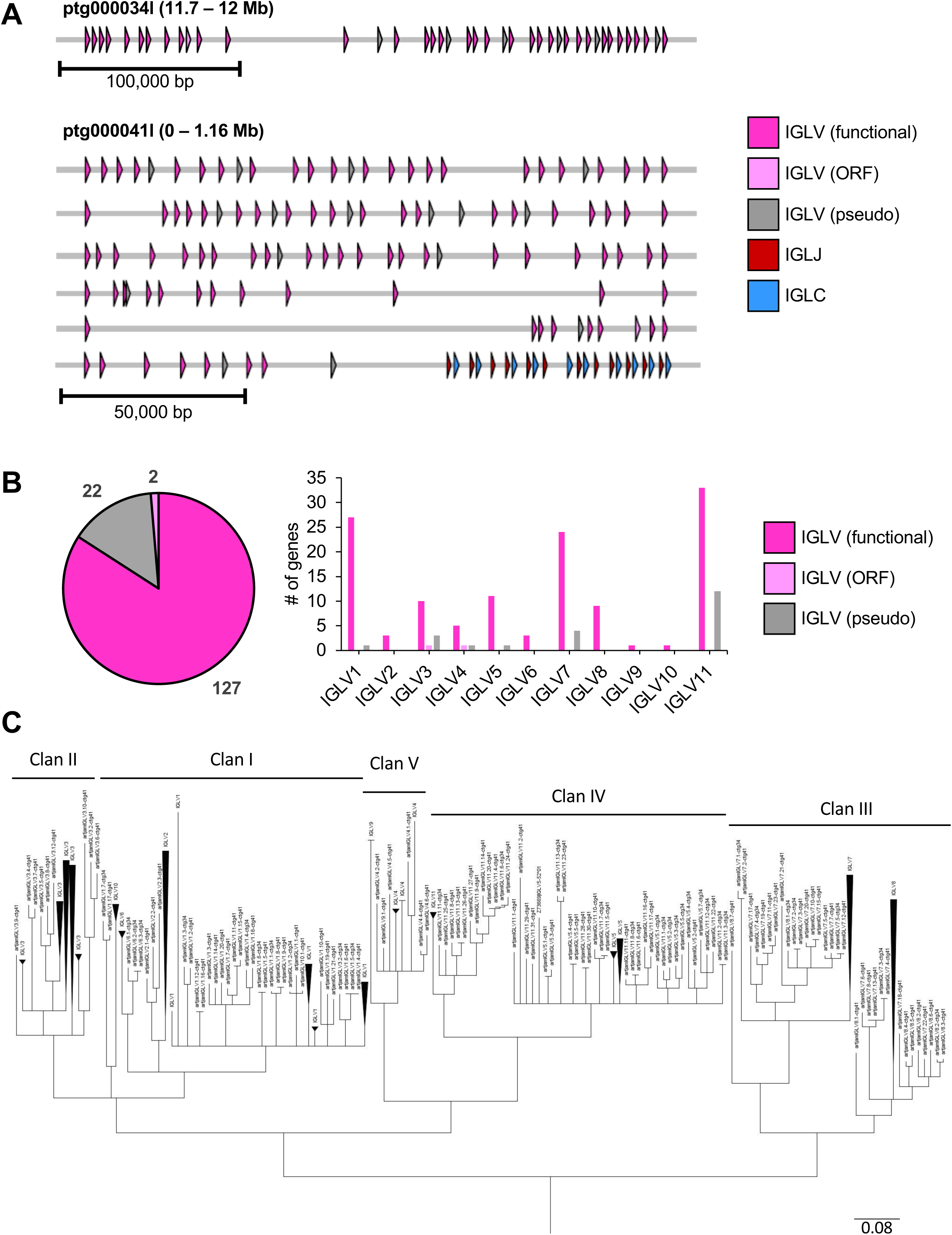
Gene organization in the *A. jamaicensis* IGL locus. **(A)** Schematic overview of the *A. jamaicensis* IGL locus on contigs ptg000034l (top) and ptg000041l (bottom). Arrows indicate directionality of the annotated genes. **(B)** Number of total functional, open reading frame (ORF), and pseudo IGLV genes annotated (pie chart) and within each IGLV family (bar chart). **©** Phylogenetic tree of all functional *A. jamaicensis* IGLV genes (n=127). Human IGLV genes (functional, pseudo, and open reading frames; without artjam prefix) are included for comparison. Clustering into IGLV clans is indicated above the tree.

We also annotated 12 IGLJ genes (11 functional, 1 pseudo) and 10 IGLC genes (all functional) (**Table S1**). In humans, IGLJ and IGLC genes are structured in a series of cassettes containing groups of IGLJ genes followed by an IGLC gene. This organization was maintained in the *A. jamaicensis* IGL locus (**Figure 3A**). To gain insight into the functionality of IGLC genes, we aligned annotated *A. jamaicensis* IGLC genes with those reported for *E. fuscus* ^16^ and humans ^28^ (**Figure S4**). Both across and within species, IGLC genes had high sequence identity (**Figure S4**). All IGLC sequences from *A. jamaicensis* and *E. fuscus* contained a single N-linked glycosylation site that was not present in human sequences (**Figure S4**). How this impacts Ig functionality will require more in-depth functional characterization.

### Immunoglobulin regulatory regions are broadly conserved

Several regulatory regions facilitate expression of IGHV genes. Here, we chose to focus on the RSS and leader sequences, as these regions are well-defined and are required for V gene expression. RSSs are conserved regions of DNA that facilitate recombination between V, D, and J genes during B cell development. They are composed of highly conserved heptamer sequences, spacers of either 12 or 23 nt, and conserved nonamer sequences ^40,41^. The RSSs associated with IGHV, IGHD, and IGHJ genes in *A. jamaicensis* were similar to those reported for both *E. fuscus* and *R. aegyptiacus* (**Figure S5A-D**) as well as those reported in humans. RSSs associated with IGLV and IGLJ genes were also similar to human and *E. fuscus* RSSs (**Figure S5E**).

The leader sequence, encoded by two exons (L-PART1 and L-PART2) directly preceding the V gene sequence, serves as the signal peptide for antibodies and has an important role facilitating *in vivo* antibody expression ^42,43^. The majority of *A. jamaicensis* leader sequences were 19 aa in length (80%), consistent with what has been reported for humans (**Table S1**) ^43^. As has been described in humans, *A. jamaicensis* leader sequences from the same IGHV family were identical or highly similar (**Table S1**) ^44^. We also compared *A. jamaicensis* IGHV leader sequences to those identified in *E. fuscus* and *P. alecto* ^16,17^. While there were differences between these three species, sequences were largely similar in length and in the types of amino acids used at each position, leading to sequences with similar physicochemical properties (**Figure S6A**). Several residues varied little between the species and across IGHV families, suggesting a potential functional importance for these residues in the expression of IGHV genes (**Figure S6A**). The majority of IGLV leader sequences were 19 aa in length (79.5%). As observed for IGHV genes, IGLV leaders from genes of the same family showed high similarity (**Table S1**). In terms of physicochemical properties, IGLV leader sequences resembled those identified for *A. jamaicensis* IGHV genes and those identified for *E. fuscus* IGLV genes ^16^, with the sequences dominated by hydrophobic and polar residues (**Figure S6B**).

Together, these data demonstrate that IGH and IGL RSS and leader sequences are broadly conserved across species, suggesting that the processes and machinery responsible for immunoglobulin gene recombination and expression are conserved as well.

### Species-specific IGH references dramatically improve BCR-repertoire analysis

Several programs exist to facilitate the analysis of BCR repertoire sequencing data, including IgBlast, IMGT/HighVQuest, and MiXCR ^45–48^. All of these tools require reference sequences to identify V, D, J, and C genes. Since few IGH references are available for bat species, analysis of BCR repertoires is typically done using references from other species; however, the complex and highly variable nature of immunoglobulin genes means that germline reference sequences can have significant impacts on estimations of BCR characteristics like somatic hypermutation (SHM) and heavy chain complementarity determining region 3 (CDRH3) length. To assess the influence of IGH references from homologous and non-homologous species on BCR repertoire analysis, we generated bulk BCR repertoire data for two adult *A. jamaicensis* infected with rabies virus (RABV). Pre-processed sequencing data (see **Methods**) were input into IgBlast using either a human reference (from IMGT ^28^), an *E. fuscus* reference (from Pursell *et al.* ^16^), or an *A. jamaicensis* reference (from our genome annotation). We then filtered the IgBlast output, retaining unique sequences with an identified IGHV, IGHD, IGHJ, and IGHC gene.

Analyses using human references performed the worst, only assigning IGHV, IGHD, IGHJ, and IGHC gene identities to about 20% of unique sequences (**Table S2**). By comparison, analyses with bat references (*E. fuscus* and *A. jamaicensis*) assigned gene identities to 50-56% of unique sequences (**Table S2**). Additionally, only analyses with bat references correctly identified IGHC genes. BCR sequencing was done using primers targeting IGHM and IGHG sequences, meaning only these isotypes were present in the datasets. However, analyses using human references incorrectly identified large numbers of sequences as IGHE or were unable to assign an IGHC identity (**Table S2**). Using bat references resulted in correct identification of IGHM and IGHG sequences, with low numbers of sequences without an IGHC identity. In this respect, analyses using *A. jamaicensis* references performed the best, with 99% of filtered sequences identified as IGHM or IGHG (**Table S2**).

We next assessed basic characteristics of the BCR repertoires to determine how reference species impacted analysis of expressed BCR repertoire characteristics. To expand our analysis, we included IGHM and IGHG BCR repertoire datasets from two uninfected *A. jamaicensis* bats from a recently published study on B cell responses to infection ^49^. Despite differences in the number of usable sequences obtained from analyses using the three reference databases, all analyses identified IGHV genes with similar frequencies across samples and isotypes (**Figure 4A-B**). This was particularly true for IGHV gene families with high levels of usage (i.e. IGHV3) (**Figure 4A-B**).

**Figure 4.**
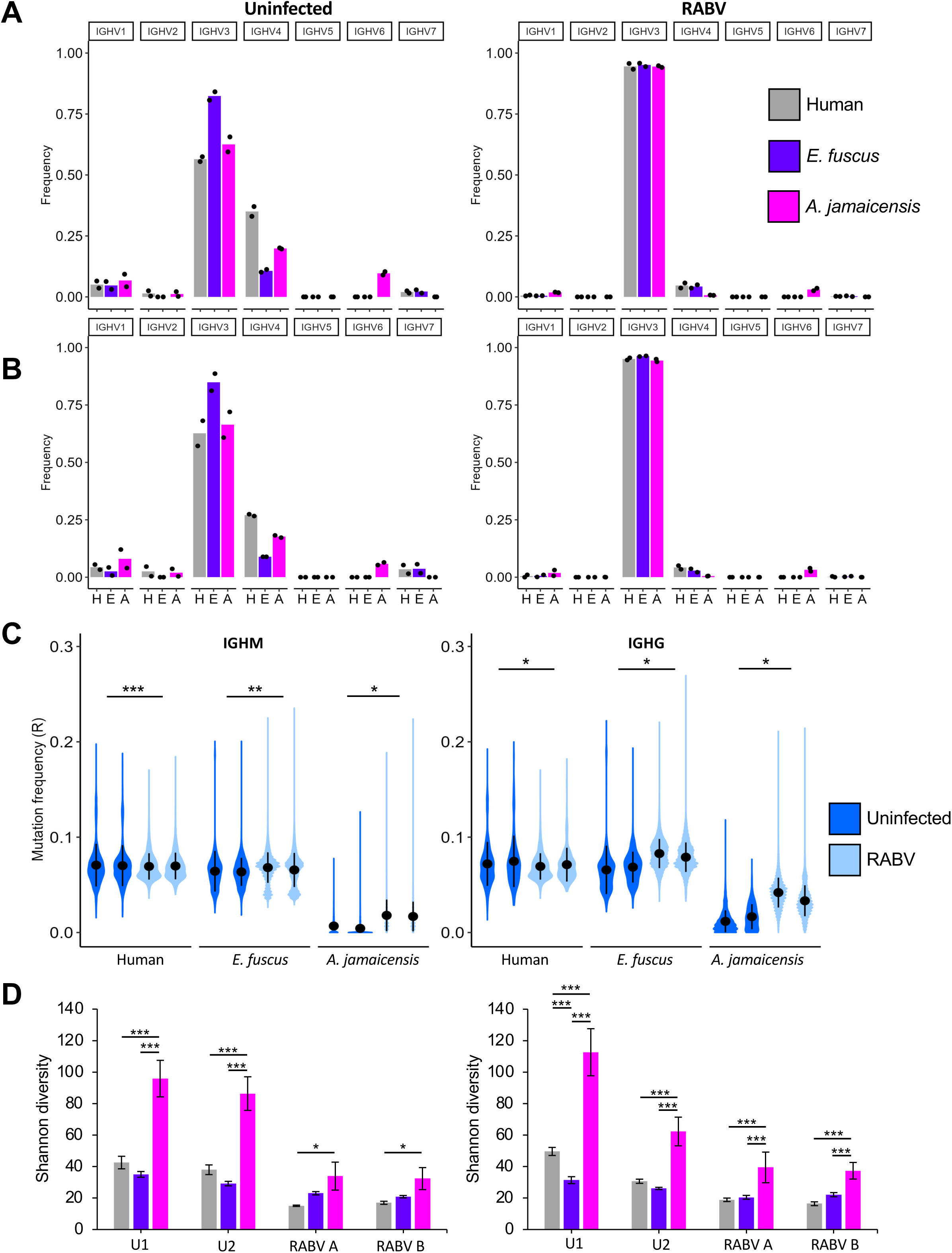
Comparison of human, *E. fuscus*, and *A. jamaicensis* IgBlast references in BCR repertoire analysis. **(A-B)** Frequency of IGHV gene families identified in **(A)** IGHM or **(B)** IGHG BCR sequences when analyzed using a human (H), an *E. fuscus* (E), or an *A. jamaicensis* (A) IgBlast reference in uninfected (n=2; from Crowley *et al.*^49^) and RABV-infected (n=2) bats. **(C)** SHM replacement (R) frequency as estimated by IgBlast with each reference in uninfected (n=2) or RABV-infected (n=2) bats. **(D)** Shannon diversity values for IGHM and IGHG BCR repertoires processed with a human, *E. fuscus*, or *A. jamaicensis* IgBlast reference for uninfected (U; n=2) and RABV-infected (RABV; n=2) *A. jamaicensis*. For this analysis, individual repertoires were sub-sampled with replacement to 1,000 sequences and repertoires processed with the same IgBlast reference database were combined. Sub-sampling was performed five times and diversity analysis was performed for each sub-sampling iteration. Higher Shannon diversity values indicate greater diversity within the sample. Bars represent the mean Shannon diversity from the five sub-sampling and diversity calculation iterations shown. Error bars represent standard deviation. Significance determined by one-way ANOVA with pairwise comparisons using Tukey’s test: *, p<0.05; ***, p<0.001.

Estimations of SHM changed based on which reference database was used to process BCR sequencing datasets. Notably, use of the human and *E. fuscus* references appeared to overestimate levels of SHM (**Figure 4C**), likely due to differences between the human and *E. fuscus* germline IGHV gene sequences and those of *A. jamaicensis*, resulting in artificially high SHM levels. Using human and *E. fuscus* references also obscured changes in SHM between healthy and infected bats. Analyses using human and *E. fuscus* references showed a negligible change in SHM in IGHM sequences in bats infected with RABV, while analyses using the *A. jamaicensis* reference captured a larger change. IGHG sequences analyzed with the human reference showed a slight *decrease* in SHM in bats infected with RABV. Sequences analyzed with *E. fuscus* and *A. jamaicensis* references captured an increase in SHM; however, analysis with the *E. fuscus* reference underestimated the magnitude of the shift, likely due to overall higher inferred levels of SHM. Across all references, differences between uninfected and RABV-infected bats were statistically significant, highlighting the importance of species-specific references for identifying the correct trend in SHM across infection conditions.

Reference databases also impacted estimations of CDRH3 length across BCR repertoires. In IGHM sequences, use of the human reference predicted the shortest average CDRH3 lengths in both uninfected and RABV-infected bats (**Figure S7A-B**). Analyses using the *E. fuscus* references calculated an identical average CDRH3 length as those using the *A. jamaicensis* reference in uninfected bats but a shorter average CDRH3 length in RABV-infected bats (**Figure S7A-B**). Regardless of infection history, use of human and *E. fuscus* references resulted in shorter calculated IGHG CDRH3 lengths than those calculated using an *A. jamaicensis* reference (**Figure S7C-D**). Differences in calculated CDRH3 length between the three references used were small, however, even small differences could mask the effects of infection on CDRH3 length.

We also investigated how the reference species influenced estimations of BCR repertoire diversity via Hill’s diversity metrics. Hill’s diversity uses input values (q) to generate an index of diversity (^q^D), with interpretable outputs at specific q values: q = 0 reflects species richness (the total number of unique BCR sequences), q = 1 reflects the Shannon diversity index where higher values indicate a more diverse and even community, and q = 2 is a transform of the Simpson diversity measurement with high values indicating the presence of a few abundant species ^50^. IGHM and IGHG BCR repertoires from both uninfected and RABV-infected bats processed with an *A. jamaicensis* reference had higher Shannon diversity values than the same repertoires processed with human or *E. fuscus* references (**Figure 4D**). While all three references captured significant shifts in IGHM and IGHG repertoire diversity between uninfected and RABV-infected bats, the magnitude of the change was largest in repertoires processed with *A. jamaicensis* references (**Figure 4D**).

### *A. jamaicensis* BCRs undergo less SHM and have shorter CDRH3s than humans

Using the captive uninfected and RABV-infected *A. jamaicensis* samples, as well as a wild-caught *A. jamaicensis* sample, we assessed the basic characteristics of the expressed BCR repertoire: IGHV gene usage, SHM frequency, and CDRH3 length to better understand the characteristics of bat B cells and determine the efficacy of our reference in the analysis of a distantly related conspecific individual (i.e. how well does a reference from a captive colony bat characterize a wild bat from Costa Rica). For both captive and wild-caught samples, pre-processed IGH repertoire sequencing data (see **Methods**) were analyzed using IgBlast and our captive *A. jamaicensis* reference ^46^.

First, we assessed the usage of IGHV and IGHJ genes in the expressed BCR repertoire of uninfected, RABV-infected, and wild-caught *A. jamaicensis*. Most germline functional IGHV genes were identified in the IgBlast data (**Table S3**). Similar patterns of IGHV gene usage were observed across IGHM and IGHG isotypes (**Figure 5A, Table S3**). In the expressed repertoire, IGHV3 and IGHV4 genes were the most commonly used genes in *A. jamaicensis* regardless of infection history or BCR isotype (**Figure 5A**). RABV infection appeared to drive an increase in the usage of IGHV3/IGHJ1 and IGHV3/IGHJ4 gene pairs (**Figure 5A**). In contrast, the wild-caught repertoires had a greater distribution of IGHV/IGHJ gene pair usage than repertoires from captive bats (**Figure 5A**). Differences in IGHV/IGHJ gene pair usage between the uninfected bats and the bats with infection history (RABV and wild-caught bats) may demonstrate evidence of selection on the expressed repertoire and are likely related to infection history as some pathogens are known to promote the usage of specific IGHV genes ^51–56^.

**Figure 5.**
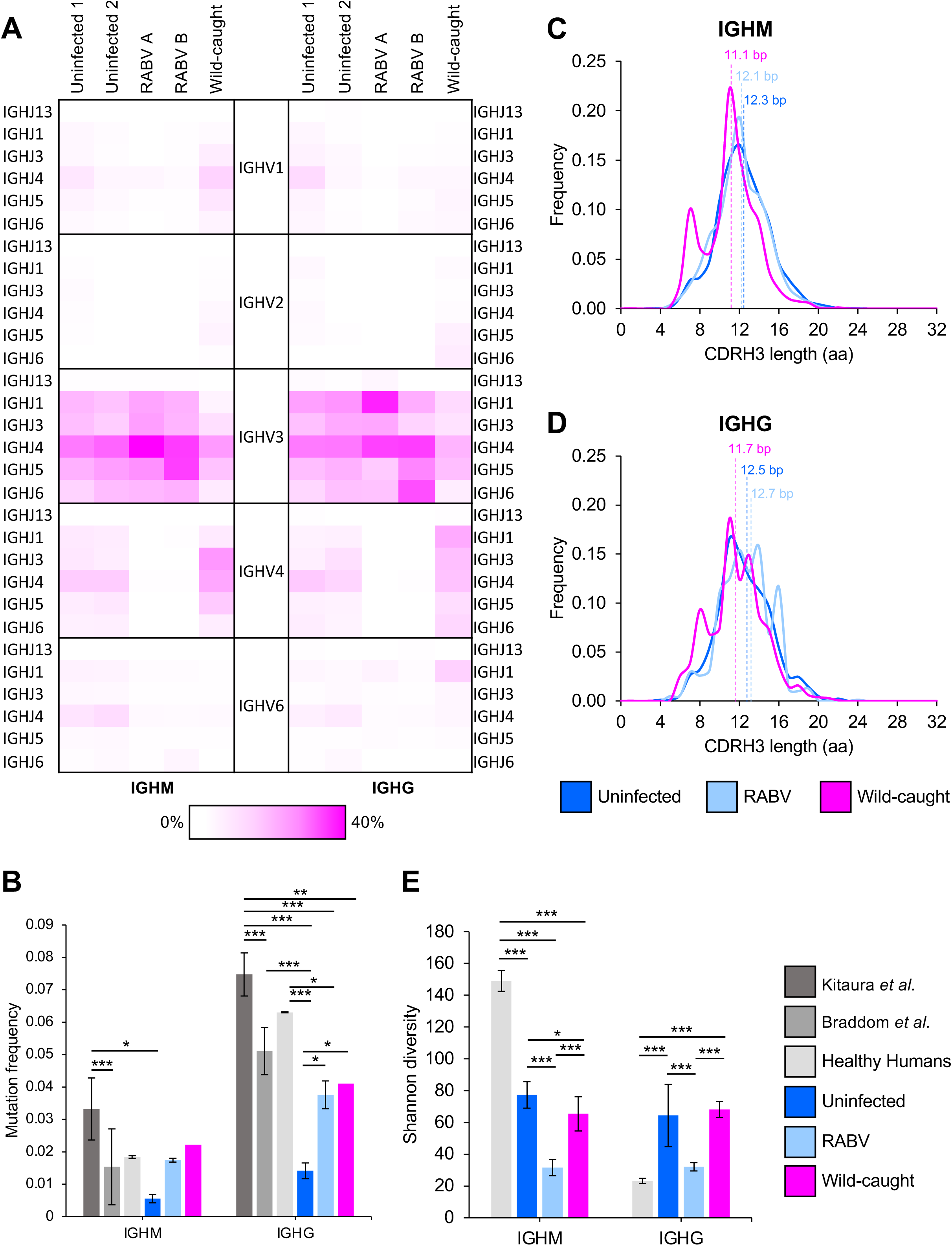
Characteristics of the B cell receptor repertoire in captive and wild-caught *A. jamaicensis*. **(A)** IGHV and IGHJ gene usage in the IGHM (left) and IGHG (right) repertoires of uninfected (n=2; from Crowley *et al.*^49^), RABV-infected (n=2), and wild-caught (n=1) *A. jamaicensis*. Cells are shaded to indicate usage frequency of the IGHV and IGHJ gene pairs. **(B)** Comparison of human and *A. jamaicensis* replacement (R) SHM frequency. Bars represent mean R SHM frequency with standard deviation plotted for each sample type (uninfected, n=2; RABV, n=2; wild-caught, n=1). Human samples were obtained from Kitaura *et al.*^57^, Braddom *et al.*^57,58^, and the Sequence Read Archive (SRA). Samples obtained from the SRA were processed in the same manner as *A. jamaicensis* samples (see **Methods**). Significant differences between mutation frequency determined by one-way ANOVA with pairwise comparisons using Tukey’s test: *, p<=0.05; **, p<0.01; ***, p<0.001. **(C-D)** Histograms of the mean frequency of CDRH3 lengths in **(C)** IGHM and **(D)** IGHG sequences from uninfected (n=2), RABV (n=2), and wild-caught (n=1) bats. The mean length for each isotype is indicated by dashed lines. **(E)** Shannon diversity of IGHM and IGHG BCR repertoires from humans (n=2; obtained from SRR7848875, SRR7849130, SRR7849308, SRR7849327) and *A. jamaicensis* (uninfected, n=2; RABV-infected, n=2; wild-caught, n=1). For this analysis, individual repertoires were sub-sampled with replacement to 1,000 sequences. Sub-sampling was performed five times and diversity analysis was performed for each sub-sampling iteration. Bars represent the mean Shannon diversity from the five sub-sampling and diversity calculation iterations. Error bars represent standard deviation. Significance determined by one-way ANOVA with pairwise comparisons using Tukey’s test: ***, p<0.001.

To gain insight into the process of SHM in *A. jamaicensis* B cells, we assessed the levels of replacement (R) SHM across the entire IGHV gene in IGHM and IGHG, BCRs from uninfected, RABV-infected, and wild-caught bats and compared them to level of R SHM observed in humans. Human samples were obtained Kitura *et al.* ^57^, Braddom *et al.* ^57,58^, and the Sequence Read Archive (Healthy Humans: SRR7848875, SRR7849130, SRR7849308, SRR7849327). The lowest levels of SHM were observed in IGHM sequences, in keeping with this population of cells undergoing less affinity maturation (**Figure 5B**) ^57,58^. *A. jamaicensis* IGHM sequences had generally comparable levels of SHM to those observed in humans, especially those from bats with a history of infection. *A. jamaicensis* IGHG sequences had significantly lower levels of SHM compared to some human samples (Kitaura *et al.*), although levels were comparable between *A. jamaicensis* and other human samples (Braddom *et al.* and Healthy Humans) (**Figure 5B**) ^57,58^. Uninfected *A. jamaicensis* IGHM and IGHG sequences had the lowest levels of SHM compared to all other samples. In IGHG sequences, SHM was significantly increased in bats that had experienced an infection (RABV and wild-caught) compared to uninfected bats (**Figure 5B**). SHM appears to increase following infection, though not necessarily to the same levels as observed in humans. Overall, these results suggest that, as has been proposed in other studies, bats rely less on SHM than humans ^5^.

Antibodies bind to their target antigens through interactions with the highly-diversified CDRs. Compared to CDRH1 and CDRH2 (encoded by the IGHV gene), CDRH3 (generated through recombination) is more varied between antibodies with differences in length and amino acid composition. The length of the CDRH3 region affects the ability of the BCR to recognize certain antigens, with longer CDRH3 sequences associated with enhanced autoreactivity and polyreactivity ^59–63^. In the expressed *A. jamaicensis* BCR repertoire, IGHM sequences had shorter CDRH3s than IGHG sequences regardless of infection history (**Figure 5C-D**). IGHM and IGHG sequences from the wild-caught bat were shorter than those observed in the captive groups. Additionally, RABV infection appeared to promote shorter CDRH3s in IGHM BCRs, but longer ones in IGHG BCRs (**Figure 5C-D**). In contrast to what we observed in *A. jamaicensis*, human naïve IGHM B cells tend to have longer CDRH3s than class-switched memory populations ^64^. In our *A. jamaicensis* data, we were unable to differentiate naïve B cells from class-switched populations by surface markers, as is typical in human studies. However, by subsetting the IGHM population based on replacement SHM frequency, we approximated a naïve B cell population. Using a mutation frequency cutoff of 0.01, we divided the IGHM B cell population into a subset with low SHM (SHM frequency < 0.01) and a subset with high SHM (SHM frequency ≥ 0.01) (**Figure S8A**) and determined the CDRH3 length for these two groups. Across all samples, BCRs with low SHM had longer CDRH3 sequences than those with higher levels of SHM (uninfected 12.4 aa vs. 11.9 aa; RABV-infected 12.3 aa vs. 12.0 aa; wild-caught 11.7 aa vs. 10.8 aa) (**Figure S8B-C**). CDRH3 lengths in IGHM B cells with low SHM were still shorter than those expressed as IGHG. This may reflect conditions in the *A. jamaicensis* immune response that favor longer CDRH3s in IGHG B cells.

Overall, *A. jamaicensis* IGHM BCR repertoires were less diverse than human repertories, regardless of infection history (**Figure 5E**). Conversely, *A. jamaicensis* IGHG repertoires were more diverse than human repertoires (**Figure 5E**). This result differs from a recent report of bat BCR diversity in *A. jamaicensis* which found that bat IGHM and IGHG BCR repertoires were more diverse than mouse repertoires following immune stimulation with a T-dependent (NP-GGG) or T-independent (NP-Ficoll) antigen ^49^. Reasons for differences between this study and our own are unclear; however, differences in sequencing depth, differences in repertoire diversity between humans and mice, or individual variation in bats could contribute. Across both isotypes, RABV infection appeared to promote less diversity (**Figure 5E**), likely due to the expansion of RABV specific clones. Within bats, IGHG diversity was high in the wild-caught sample (**Figure 5E**), likely reflecting diversification of the BCR repertoire in response to multiple pathogen exposures.

### Cysteine-rich IGHV genes are enriched in *A. jamaicensis*

Structurally, cysteine residues have critical roles, forming intra- and inter-chain disulfide bonds within antibodies ^65^. All functional IGHV genes encode two canonical cysteines, one at the end of FWR1 and one at the end of FWR3; these residues form a disulfide bond that is essential for the three dimensional structure of the antibody variable region ^14^. Non-canonical cysteines in other regions of the antibody have been identified in numerous species, including humans, and contribute to antibody repertoire diversity ^66–70^. We found several IGHV genes in *A. jamaicensis* with up to four additional, non-canonical cysteines. In total, 26% (15/57) of functional IGHV genes had three or more total cysteines. Using the amino acid sequences of functional IGHV genes available from IMGT and a recent *E. fuscus* IGH annotation ^16,28^, we compared the number of functional genes with three or more total cysteine residues (referred to as cysteine-rich) in eight species to the number found in *A. jamaicensis* (**Figure 6A**). Generally, levels of cysteine-rich IGHV genes were low (4%-11%) across non-bat species. In humans, only 7% (21/323) of functional IGHV gene alleles were cysteine-rich. The only species with comparable levels of cysteine-rich IGHV genes to *A. jamaicensis* were *E. fuscus* (32%, 24/76) and cows (*Bos taurus*; 40%, 10/25). However, *A. jamaicensis* and *E. fuscus* were the only species with IGHV genes that contained five or six total cysteines, representing a potentially bat-specific expansion of cysteine-rich IGHV genes. All but one of the non-canonical cysteine residues in *A. jamaicensis* fell at the end of CDR1, within FWR2, or at the beginning of CDR2 (**Figure 6B**). Cysteines in these regions may be able to form disulfide bonds between CDR1 and CDR2 or with CDR3.

**Figure 6.**
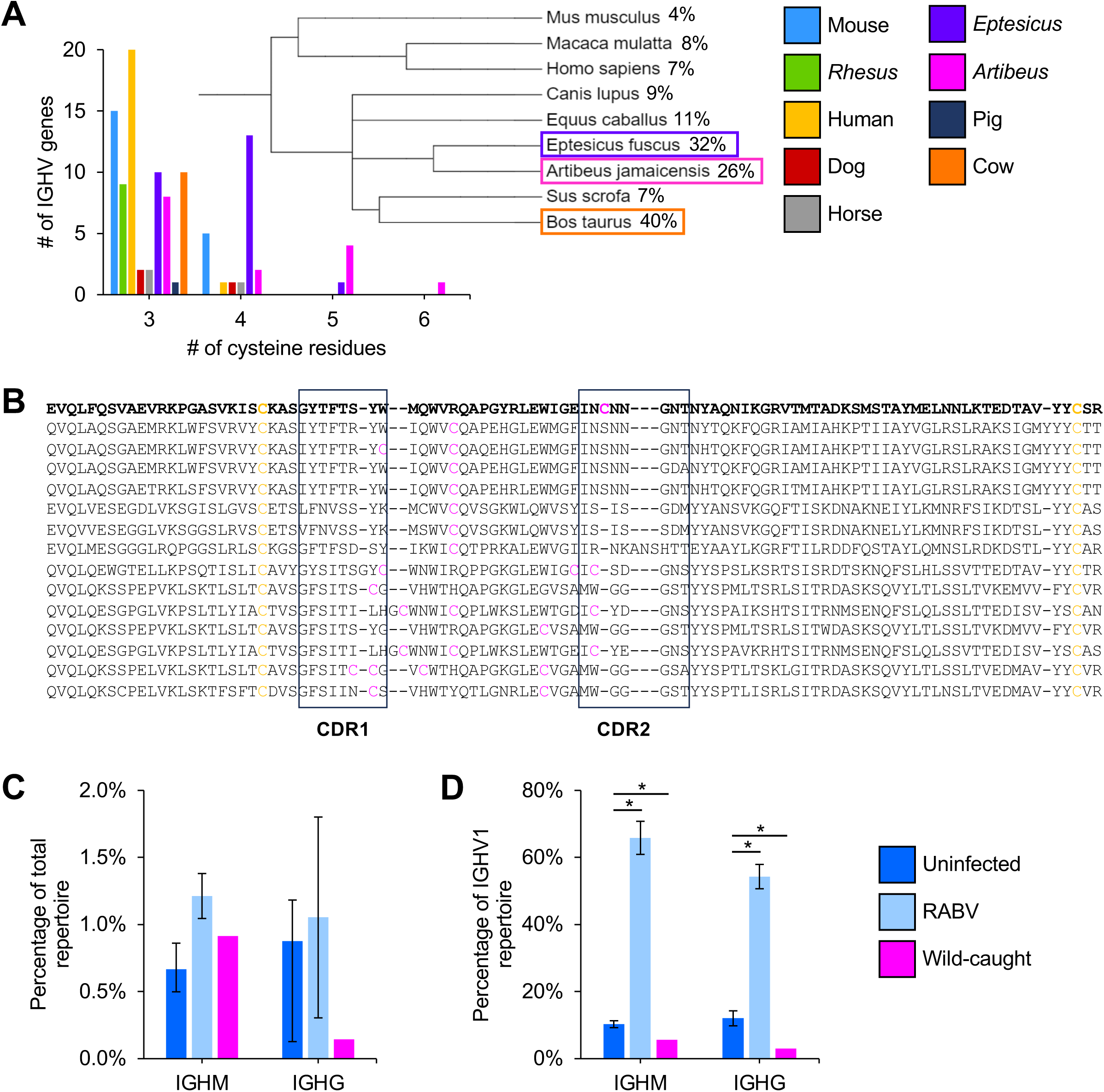
Cysteine-rich IGHV genes in *A. jamaicensis*. **(A)** Number of genes with 3 or more cysteine residues across species (bar chart) with species relationships depicted in the phylogenetic tree. Percentage of IGHV genes with 3 or more cysteine residues are displayed next to the species names. **(B)** Amino acid sequence alignment of the 15 functional *A. jamaicensis* IGHV genes with 3 or more cysteine residues. Canonical cysteines are shown in yellow. Additional, non-canonical cysteines are shown in pink, CDR1 and CDR2 regions are denoted by rectangles, and artjamIGHV1.23-ctg26 is bolded. **(C)** Frequency of IGHM and IGHG sequences using artjamIGHV1.23-ctg26 in the total repertoire of uninfected (n=2), RABV-infected (n=2), and wild-caught (n=1) bats. No comparisons were statistically significant by one-way ANOVA. **(D)** Frequency of IGHM and IGHG sequences using artjamIGHV1.23-ctg26 in the IGHV1 repertoire of uninfected (n=2), RABV-infected (n=2), and wild-caught (n=1) bats. *, P < 0.05 as determined by one-way ANOVA with pairwise comparisons using Tukey’s test. In plots C-E, bars represent the mean and error bars represent the standard deviation. Bars represent the mean difference of SHM frequency in uninfected (n=2), RABV-infected (n=2), and wild-caught (n=1) bats. Error bars indicate the standard deviation. *, p<0.05 as determined by one-way ANOVA with pairwise comparisons using Tukey’s test.

While there was an expansion of cysteine-rich IGHV genes in the *A. jamaicensis* germline, they were expressed at low levels. Only a single cysteine-rich IGHV gene, artjamIGHV1.23-ctg26, was detected in the uninfected, RABV-infected, and wild-caught individuals. Expression of this gene was low in IGHM (0.5-1.4% across all samples) and IGHG (0.3-1.8% across all samples) repertoires (**Figure 6C, Table S3**). Expression of this gene may increase in response to RABV infection, however high variability between individuals makes it difficult to draw a clear conclusion. Interestingly, in RABV-infected bats, artjamIGHV1.23-ctg26 accounted for the majority of the expressed IGHV1 repertoire (**Figure 6D**).

Determining the roles these cysteine-rich IGHV genes play in B cell responses in bats will require further investigation, potentially using approaches that specifically target these genes.

## Discussion

Immunoglobulins are critical to vertebrate immunity and derive much of their diversity from complex genetic loci and somatic mutation. This complexity complicates efforts to characterize Ig loci outside of model organisms. Accordingly, few studies have characterized immunoglobulin loci in bats, often superficially (e.g. IGH in *R. ferrumequinum, P. discolor, and P. pipistrellus*), or even failing to note an important IGH duplication (*P. pipistrellus*) ^15^ (but see ^16^). Additionally, only one study has annotated the bat IGL locus (*E. fuscus*) ^16^. Inaccuracies in previous studies demonstrate the difficulties involved in sequencing and characterizing this locus, further highlighting the need for increased attention in this area. Accurate references are crucial for understanding more complicated aspects of the expressed repertoire, like SHM frequency. Here, we generated a detailed, high-quality germline reference for the *A. jamaicensis* immunoglobulin loci and used it to characterize the basic features of the BCR repertoire of this species, while demonstrating the importance of species-specific reference databases for BCR repertoire characterization.

Generally, *A. jamaicensis* maintained the structure and composition of the IGH and IGL loci that has been observed in humans and other mammals, including the five bat species for which IGH annotations are available ^2,15,16^. However, even within this conserved locus, we have observed notable differences between bat species, particularly in the number and types of IGHV genes (*A. jamaicensis*, 196 IGHV; *E. fuscus*, 132 IGHV; *R. aegyptiacus*, 66 IGHV), demonstrating that while aspects of adaptive immunity may be largely similar across bats, there are species-specific differences. This supports findings from studies of innate immunity indicating that bats are not a monolith ^22,71,72^. The functional impact of species-specific differences in the composition of the IGH locus remain unclear, but they provide a potential clue to determining the underlying drivers of high variability in humoral responses observed between species.

While we present one of the most detailed analyses of bat immunoglobulins to date, this study is not without limitations. Annotations and analyses were done with human-centric references and programs which may bias our results to highlight more conserved sequences, potentially masking unique, bat-specific features of the IGH and IGL loci that are more divergent from humans. In particular, RSS sites were identified based on similarity to canonical human sequences, leaving the possibility that there are other non-canonical RSSs in bats. More in-depth BCR sequencing of expressed repertoires may help uncover functional V genes in the germline that use RSS sites not identified in this study.

Here, we have generated a valuable tool for the bat research community in the form of a highly-contiguous, annotated genome with detailed IGH and IGL locus annotations. To continue making meaningful advancements in the area of bat humoral immunity, we need to prioritize studies that provide insight into B cell characteristics and generate tools for deeper understanding of the adaptive immune response. Bat research has long suffered from a lack of tools in comparison to human and mouse studies. Developing the basic tools we have for humans and mice, like germline genome references, for bat species will facilitate deeper investigations into bat humoral immunity.

## Resource availability

### Lead contact

Further information and requests for resources and reagents should be directed to and will be fulfilled by the lead contact, Hannah Frank.

### Materials availability

This study did not generate new unique reagents.

### Data code and availability

The *A. jamaicensis* genome assembly and annotation is available at GenBank (PRJNA1244911). BCR repertoire datasets generated as part of this study have been deposited at SRA (PRJNA1244374) and are publicly available as of the date of publication.

All original code is available upon request.

## Acknowledgements

We thank Yana Safonova and Anton Bankevich for their assistance with the genome assembly strategy. We thank Scott Boyd and his laboratory for his support and guidance in obtaining the *A. jamaicensis* repertoire sequencing. We gratefully acknowledge the University of Louisville Sequencing Technology Center who performed the PacBio HiFi sequencing, and the Organization for Tropical Studies, the Las Cruces Biological Station and Jon Flanders for support during fieldwork. This study was funded by NIH NIAID (R21 AI169548 to H.K.F. and T. S.), NIH NIAID R24AI165424 (T.S.), Life Sciences Research Foundation Fellowship (H.K.F.), Open Philanthropy Project, Stanford Woods Environmental Venture Program, Bing-Mooney Fellowship in Environmental Science (H.K.F.), Stanford Center for Computational, Evolutionary and Human Genomics Postdoctoral Fellowship (H.K.F.), NIH (5 T32 AI007290, H.K.F.), and Stanford School of Medicine Dean’s Postdoctoral Fellowship (H.K.F.).

## Author contributions

H.K.F., T.P., and A.B.R. conceptualized the study. H.K.F. sampled and generated the expressed BCR repertoire from the wild-caught *A. jamaicensis.* T.S. provided the *A. jamaicensis* cells used for genome sequencing. S.Z., C.R., N.H., and T.S. performed RABV infections and sample collections from *A. jamaicensis* bats. A.B.R. assembled the *A. jamaicensis* genome, annotated the IGH and IGL loci, prepared BCR sequencing libraries from captive *A. jamaicensis* bats, and analyzed the expressed BCR repertoire data. T.P. sequenced the BCR repertoire libraries from captive RABV-infected bats. T.L. performed the TOGA whole-genome annotation. A.B.R. and H.K.F. wrote the manuscript. All authors edited, read, and approved the final manuscript.

## Declaration of interests

The authors declare no competing interests.

## Supplementary information

Document S1. Figures S1-S9

Table S1. Excel file containing information about genes annotated in the IGH and IGL loci, related to Figures 2 and 3

Table S2. Excel file containing BCR repertoire sequencing and IgBlast statistics, related to Figures 4, 5, and 6

Table S3. Excel file containing information about IGHV gene expression across all samples, related to Figure 5

## Supplemental Figure Legends

Figure S1. Distribution of the immunoglobulin locus across contigs in two *Artibeus jamaicensis* genome assemblies, related to Figure 1. IgDetective output showing the distribution of IGH, IGK, and IGL genes across contigs from **(A)** CSHL_Jam ^22^ and **(B)** our PacBio genome assemblies. Numbers in shaded squares indicate the number of predicted genes on each contig.

**Figure S2. IGHC gene alignments, related to Figure 2**. Alignment of **(A)** IGHM, **(B)** IGHG, **(C)** IGHE, and **(D)** IGHA amino acid sequences from *A. jamaicensis* (ArtJam), *E. fuscus* (EptFus) (from the IGH locus on both chromosome 5 and chromosome 24), and humans. In all alignments, cysteine residues are in pink and N-glycosylation sites are indicated by bold and underlined text. N-glycosylation sites unique to *A. jamaicensis* are highlighted yellow, while sites present in bats (*A. jamaicensis* and *E. fuscus*), but not humans, are highlighted green. *A. jamaicensis* sequences labeled “transcript” are those sequences lacking an M domain, while those labeled “M” contain the M domains. **(B)** FcγR1 binding sites in the CH2 region (Leu-Leu-Gly-Gly in the lower hinge region, Asp^265^ in the CH2 BC loop, N-glycosylation site at Asn^297^, and Ala-Leu-Pro-Ala-Pro in the CH2 FG loop^31–33^) are in green. The Cys-X-X-Cys motif conserved in human IGHG hinge regions^73^ is shown in purple. **(C)** FcεR1 binding sites in the CH3 region (Ser-Arg-Ala-Ser-Gly-Lys-Pro-Val-Asn-His-Ser and Arg-Ala-Leu-Met^74^) are in blue. **(D)** Domains in the CH2 (Leu-Leu) and CH3 (Pro-Leu-Ala-Phe) regions implicated in IGHA1 binding to FcαR^75^ are in orange. In all panels, individual exons are indicated by colored bars under the alignments: CH1, blue; H, grey; CH2, purple; CH3, green; CH4, pink; M1 and M2, yellow.

**Figure S3. IGHC gene expression and functionality, related to Figure 2. (A)** Switch motif density plotted across the genome region containing IGHC genes. The frequency of AID motifs (AGCT) on the forward and reverse strand per 1,000 bp was used to calculate Z-scores. Relative location in the locus is depicted on the x-axis (bp) and boxes under the plot indicate the location of the IGHM, IGHG, IGHE, and IGHA genes. Red arrows indicate areas of increased switch motif density preceding IGHC genes. **(B)** BLAST alignment of RNA-seq reads from an *A. jamaicensis* spleen to the indicated coding regions of IGHM, IGHG, IGHE, and IGHA genes annotated in our PacBio genome. Up to 100 sequences with significant alignments are depicted below the schematic diagram of IGHC coding regions. The color of each sequence indicates the alignment score determined by BLAST. **(C)** Bulk tissue expression (median transcripts per million, TPM) for indicated IGHC genes in the human spleen. Data were obtained from the GTEx Portal on 07/11/2024 (dbGaP accession number phs000424.v8.p2).

**Figure S4. IGLC gene alignments, related to Figure 3**. Alignment of IGLC amino acid sequences from *A. jamaicensis* (ArtJam), *E. fuscus* (EptFus), and humans. Cysteine residues are shown in pink and N-glycosylation sites are indicated by bold and underlined text. N-glycosylation sites unique to bats (*A. jamaicensis* and *E. fuscus*), but not humans, are highlighted green.

**Figure S5. IGH and IGL RSS sequences, related to Figures 2 and 3. (A-D)** Frequency of nucleotides at each position of the RSS23 and RSS12 heptamer and nonamer sequences are shown for *A. jamaicensis* (A), *E. fuscus* (E), and *R. aegyptiacus* (R) for the **(A)** IGHV 5’ RSS23 sequence (A, n=195; E, n=120; R, n=6), **(B)** IGHJ 3’ RSS23 sequence (A; n=6; E, n=11; R, n=9), **(C)** IGHD 5’ RSS12 sequence (A, n=7; E, n=19; R, n=8), and **(D)** IGHD 3’ RSS12 sequence (A, n=7; E, n=19; R, n=8). **(E)** Frequency of nucleotides at each position of the IGLV RSS23 (right) and IGLJ RSS12 (left) heptamer and nonamer sequences are shown for *A. jamaicensis* (A; IGLV n=110, IGLJ n=11) and *E. fuscus* (E; IGLV n=126, IGLJ n=12).

**Figure S6. IGHV and IGLV leader sequences, related to Figures 2 and 3. (A)** Frequencies of amino acids at each position of the IGHV leader sequence are shown for *A. jamaicensis* (A; n=27), *E. fuscus* (E; n=107), and *P. alecto* (P; n=23). **(B)** Frequencies of amino acids at each position of the IGLV leader sequence are shown for *A. jamaicensis* (A, n=101) and *E. fuscus* (E, n=73). In all plots, only leaders with a length of 19 amino acids are shown, and sequences are colored based on physicochemical properties.

**Figure S7. Effect of reference database on CDRH3 length calculations, related to Figure 4. (A)** Histograms of the frequency of CDRH3 lengths of **(A)** IGHM sequences from uninfected bats (n=2), **(B)** IGHM sequences from RABV-infected bats (n=2), **(C)** IGHG sequences from uninfected bats (n=2), and **(D)** IGHG sequences from RABV-infected bats (n=2). The mean length for each isotype is indicated by dashed lines.

**Figure S8. Comparison of naïve-like IGHM B cells to the IGHM repertoire, related to Figure 5. (A)** Replacement (R) mutation frequency in all IGHM sequences (T), those with SHM < 0.01 (L), and those with SHM ≥ 0.01 (H) in repertoires from uninfected (n=2), RABV-infected (n=2), or wild-caught (n=1) bats. Within the boxplot, the center line indicates the median, upper and lower limits of the box indicate the first and third quartiles, and the whiskers indicate the upper and lower limits of the data within 1.5 times the interquartile range. Values outside the bounds of the whiskers are plotted individually. **(B-D)** CDRH3 length of IGHM BCR sequences with low SHM frequency (SHM < 0.01) or high SHM frequency (SHM ≥ 0.01) in **(B)** uninfected (n=2), **(C)** RABV-infected (n=2), and **(D)** wild-caught (n=1) bats. Average CDRH3 lengths for each group are indicated by dashed lines. Significant differences between the CDRH3 lengths in low SHM and high SHM groups was assessed by Student’s t-test. P-values reported on the graphs.

**Figure S9. Detection of rabies virus following infection in *A. jamaicensis* bats, related to Figures 4, 5 and 6. (A)** Gel electrophoresis showing PCR amplification of RABV (CSV-N2c) from saliva samples collected at 23 days post infection from *A. jamaicensis* infected with RABV (n=2). Amplified RABV-CSV-N2c RNA was included as a positive control. Expected band size for RABV is 796 bp. **(B)** Unedited image of the gel shown in **(A)**.

## Methods

### Genome sequencing and assembly

Primary kidney cells derived from a male *Artibeus jamaicensis* bat (AJK6) were frozen at a low passage number in 10% FBS and 10% DMSO-DMEM at a concentration of 1 million cells.

Cells were sent to University of Louisville where high molecular weight DNA extraction and PacBio HiFi library preparation was performed according to the manufacturer’s recommendations. PacBio HiFi reads were assembled using Hifiasm (v0.16.1-r375) with parameters -l2 and --primary. N50 comparisons to other genomes [*A. jamaicensis* CHSL_Jam (GCA_021234435.1), *Desmodus rotundus* (GCA_022682495.1), *Phyllostomus discolor* (GCA_004126475.3), *Myotis daubentonii* (GCA_963259705.1), *Eptesicus fuscus* (GCA_027574615.1), *Rhinolophus ferrumequinum* (GCA_004115265.3)] were calculated using Quast (v5.2.0) ^76,77^. The completeness of the assembly was assessed using an a priori set of 12,234 conserved genes using BUSCO (v5.2.2; laurasiatheria_odb10) in genome mode ^78,79^. The *A. jamaicensis* assembly contained 95.3% of genes in BUSCO’s laurasiatheria_odb10 set. The remaining 4.7% of conserved genes were classified as fragmented (N=99) or missing (N=487).

Comparison between our PacBio assembly and the CSHL_Jam assembly was accomplished with minimap2 (v2.26-r1175), and the two assemblies were aligned using -x asm20 and default settings ^80^. CHSL_Jam contigs that aligned to our PacBio contigs of interest with at least 70% identity over a region of at least 100,000 bp were extracted and alignments were visualized using the R function asynt ^81^.

### Genome annotation

#### TOGA whole-genome annotation

A *de novo* repeat library was created using RepeatModeler (v2.0.4; flag --engine NCBI) for our PacBio genome. Repeats in the genome assembly were then soft-masked with RepeatMasker (v4.1.5; flag --engine crossmatch -s). TOGA (Tool to Infer Orthologs from Genome Alignments) is a projection-based annotation tool which requires a high-quality reference annotation ^26^. Orthologs were inferred using 36,664 transcripts and 19,456 coding genes from the GENCODE v38 (Ensembl 2014) annotation of the human genome (hg38) as a reference. Local alignments generated by LASTZ were prepared as TOGA inputs with human-to-other-mammal DEF file parameters (K=2200, L=6000, Y=3400, H=2000). Chaining was performed using axtChain with customized parameters (flag --chain_linear_g=medium) and local alignments were improved using RepeatFiller and chainCleaner with default parameters (flag --minBrokenChainScore=75000). Pairwise genome alignment chains between human and *A. jamaicensis* were used to infer and annotate intact orthologous genes with TOGA. The completeness of the annotation was assessed using an a priori set of 9,226 near-universally conserved mammalian genes using BUSCO (v5.2.2; odb10_mammalia) in protein mode. The *A. jamaicensis* annotation contains 99.7% of genes in BUSCO’s odb10_mammalia set. The remaining 0.3% of conserved genes were classified as fragmented (N=4) or missing (N=23). BUSCO does not distinguish whether genes are classified as fragmented or missing due to assembly incompleteness or base errors which shift the reading frame. However, annotation completeness is highly correlated with assembly completeness calculated by BUSCO in genome mode ^26^.

#### IGH locus annotation

We identified contig(s) potentially containing the IGH locus using IgDetective (v1.1.0) with default settings on our PacBio genome assembly ^27^. This identified a single contig containing both IGHV and IGHC genes (ptg000026l) as well as an additional contig containing a single IGHV gene (ptg000032l). Next, using the Geneious (v2023.2.1) “Map to Reference” function (medium sensitivity, iterate up to 5 times), we aligned all IGHV, IGHD, IGHJ, and IGHC nucleotide sequences available from IMGT ^28^ to contig ptg000026l and ptg000032l. Using the same settings, we also aligned the predicted IGHV genes (*de novo* search, predicted genes IGH) reported by IgDetective to both contigs. The predicted IGHV gene on contig ptg000032l did not pass further annotation processes (described below).

For each region where an IMGT or IgDetective IGHV reference sequence aligned, we translated the alignment region. Based on the translations, some annotations were extended beyond the alignments to capture the full IGHV gene. If there were no stop codons in the translation, we next annotated the leader sequence (L-PART1, intron, and L-PART2). Leaders were considered functional if they contained no stop codons and had canonical donor and acceptor splice sites. For all IGHV gene alignments, regardless of stop codons in the translation, we searched for RSS23 sites at the end of the alignment using the Recombination Signal Sequence Site ^82^ and human RSS23s as a reference. RSSs were deemed functional if they had a passing RIC score (RIC > -58.45) and were oriented in the same direction as the IGHV gene. Annotations were extended when necessary, so that the RSS23 directly followed the IGHV gene with no gaps between these two sequences. Genes were deemed functional if they possessed an in-frame leader sequence with functional donor and acceptor splice sites, had no stop codons in the IGHV coding sequence, and had an RSS23 sequence immediately following the IGHV sequence. IGHV genes with an out of frame leader sequence, mutated splice sites, or stop codons in the leader or IGHV coding sequence were deemed pseudogenes. Otherwise functional IGHV genes lacking a RSS23 sequence were labeled ORFs. All identified IGHV gene sequences (nucleotide and amino acid translation) and leader sequences are provided in **Table S1**.

We identified five IGHD genes based on alignments with IMGT reference sequences. Using the Recombination Signal Sequence Site with human RSS12s as a reference, we annotated RSS12 sites on the 5’ (reverse orientation) and 3’ (forward orientation) ends of the IGHD alignments. Only RSS12s with a passing RIC score (RIC > -38.81) were considered functional. These alignments were extended as necessary so that the RSS12 sites directly preceded and followed the IGHD annotation. IGHD genes are short and can be difficult to locate by alignment alone. For this reason, to identify additional IGHD genes, we searched the region between the last IGHV gene alignment and the first IGHJ gene alignment for RSS12 sites. Sequences between two, correctly oriented RSS12 sites were annotated as probable IGHD genes. Identified IGHD gene sequences are provided in **Table S1**.

For all regions with IMGT IGHJ reference sequences aligned, we identified RSS23 sites 3’ (reverse orientation) to the alignment region using the Recombination Signal Sequence Site with human RSS23s as a reference. Only RSS23s with a passing RIC score (RIC > -58.45) were considered functional. We then translated the alignment region (extended as necessary to ensure the IGHJ gene directly followed the RSS23) in all three reading frames. The reading frame that contained the conserved IGHJ motif 5’-WGXG-3’ was considered the functional reading frame. Final IGHJ annotations were adjusted as necessary to account for splice sites at the end of the annotation necessary for splicing to IGHC genes. IGHJ sequences and translations in the functional reading frame are provided in **Table S1**.

Each IGHC gene is composed of several exons. IMGT reference sequences for IGHM, IGHD, IGHG, IGHE, and IGHA were obtained for each exon region (CH1, CH2, H (for IGHG), CH3, CH4 (for IGHM and IGHE), CHS, and M1/2), allowing us to identify the specific regions of each isotype based on the alignment of the IMGT references to our genome assembly. Regions with IMGT reference sequences aligned were adjusted to account for splice sites, when necessary, and translated to check for stop codons. Translations were also compared to human and *E. fuscus* IGHC translations using the Geneious “Multiple to Align” function (MUSCLE Alignment, PPP algorithm) to check for similarities in sequence identity, binding motifs, and length. Sequences were considered functional if they contained no stop codons and canonical splice sites.

Our alignment did not identify an IGHD gene. To further confirm the absence of IGHD in *A. jamaicensis*, we used BLAST ^83,84^ to query the genomic region between the IGHM and IGHG annotations for the human IGHD sequence. No sequence similarity was found between the *A. jamaicensis* genome region and IGHD. Sequence and translation information for the identified IGHC genes are available in **Table S1**.

#### IGL locus annotation

The *A. jamaicensis* light chain locus was annotated through the same process as described for the IGH locus annotation. In brief, to identify contig(s) potentially containing the immunoglobulin light chain lambda (IGL) and light chain kappa (IGK) loci, we utilized IgDetective (v1.1.0) with default settings and our PacBio genome ^27^. This identified a single contig containing both IGLV and IGLC genes (ptg000041l) as well as an additional contig containing IGLV genes (ptg000034l). No IGK genes were identified by IgDetective. The absence of an IGK locus was confirmed by searching our PacBio genome for IGKV and IGKC sequences with BLAST ^85,86^. No sequences with similarity to IGK genes were identified. Next, using the Geneious (v2023.2.1) “Map to Reference” function (medium sensitivity, iterate up to 5 times) we aligned all IGLV, IGLJ, and IGLC nucleotide sequences available from IMGT ^28^ to contig ptg000041l and ptg000034l. Using the same settings, we also aligned the predicted IGLV genes (*de novo* search, predicted genes IGL) reported by IgDetective to both contigs.

For each region where an IMGT or IgDetective IGLV sequence aligned, we translated the alignment region, identified RSS23 sites, extended the annotation as necessary to capture the entire IGLV sequence, and annotated a leader sequence for IGLV genes with no stop codons in the coding region. Functionality was assigned based on IMGT definitions as described for our IGH locus annotation. All identified IGLV gene sequences (nucleotide and amino acid translation) and leader sequences are provided in **Table S1**.

For all regions with IMGT IGLJ reference sequences aligned, we identified RSS12 sites 5’ (reverse orientation) to the alignment region. We then translated the alignment region in all three reading frames. The reading frame that contained the conserved IGLJ motifs 5’-FGXG-3’ and 5’-TVL-3’ was considered the functional reading frame. Annotations were adjusted to account for splice sites necessary for splicing to IGLC genes. IGLJ sequences and translations in the functional reading frame are provided in **Table S1**.

For IGLC genes, regions where IMGT IGLC reference sequences aligned were annotated in the *A. jamaicensis* genomes and translated to check for stop codons. *A. jamaicensis* IGLC translations were aligned with human and *E. fuscus* IGLC genes to check for sequence similarity using the Geneious “Multiple to Align” function (MUSCLE Alignment, PPP algorithm). IGLC sequences and translations are provided in **Table S1**. All locus schematics were created using the R package gggenes (v0.5.1) ^87^.

### Gene labeling

When labeling the IGHV genes annotated in the *A. jamaicensis* IGH locus, we chose to designate IGHV gene family identity based on the IGHV gene family of the IMGT or IgDetective sequences that aligned to the *A. jamaicensis* genome. *A. jamaicensis* IGHV gene annotations were then numbered within IGHV families starting with the gene farthest from the IGHC region, such that the first IGHV3 gene identified was labeled IGHV3.1, the second IGHV3.2, and so on. The first IGHV4 gene was labeled IGHV4.1, the second IGHV4.2, etc. Pseudogenes were labeled with “p” and open reading frames with “orf.” IGHJ genes were labeled based on the IGHJ family identity of the IMGT alignment. IGHD genes were labeled in the same fashion, however IGHD genes identified based on RSS12 sites alone were labeled as “probable IGHD” genes. IGHC genes were labeled based on the human IGHC gene alignment. IGLV and IGLJ genes were labeled as described for IGH genes. IGLC genes were labeled based on the human IGLC gene alignment.

### *Artibeus jamaicensis* samples

#### Captive bats

All captive *A. jamaicensis* bats were obtained from a breeding colony maintained at Colorado State University ^24,88^. This colony has been closed for 18 years and is rabies virus free. For RABV-infected samples, two separate adult, male *A. jamaicensis* bats were inoculated intramuscularly in the leg with 10^4^ focus forming units (FFU) CSV-N2c rabies virus while anesthetized with 3% isoflurane in O_2_. Bats were monitored for symptom development and euthanized at 23 days post infection (dpi). Terminal spleen samples were flash frozen. Presence of rabies virus in both bats was verified by PCR on saliva samples taken at 21 dpi (**Figure S9**). The research was approved by the Colorado State University Institutional Animal Care and Use Committee (protocol #4565).

#### Wild-caught bat

A single, male *A. jamaicensis* individual was captured in a mist net and euthanized in accordance with AVMA guidelines at the Las Cruces Biological Station in Costa Rica. The spleen was removed and stored in RNAlater for 24 hours at room temperature and then frozen at -20°C and later -80°C until further processing. Collection and analysis was conducted under Costa Rican permits R-044-2015-OT-CONAGEBIO and R-054-2019-OT-CONAGEBIO. The research was approved by the Stanford Institutional Care and Use Committee (protocol #29978).

### BCR-repertoire amplification

For captive, RABV-infected *A. jamaicensis* samples, RNA was extracted from splenic tissue with the Quick-RNA/DNA miniprep plus (Zymo #D7003) kit, according to the manufacturer’s instructions. cDNA was prepared using the SuperScript III First Strand Synthesis System (Invitrogen #18-080-051) according to the manufacturer’s instructions. BCR sequences were amplified from cDNA by PCR using a pool of forward primers targeting the beginning of IGHV genes (FR1 primer mix: 5’-GACGTGCAGCTAGTGGAGTCTGG-3’, 5’-GAGGTGCAGCTGGTGGAGTCT-3’, 5’-CAGGTGCAGCTTAAGGAGTCAGGTC-3’, 5’-GAAGTGCAGCTGGTGGAGTCTGG-3’, 5’-CAGTTGCAGCTGCAGGAGTCG-3’, 5’-GAAGTGCAGCTATTTCAGTCTGTGGC-3’, 5’-CAAGTGAAGCTGCAGGAGTCTGGC-3’, 5’-CAGGTCACCCTGCAGGAGTCTG-3’, 5’-CAGGTACAGCTGGTACAGTCTGGG-3’) and a reverse primer targeting the CH1 region of either IGHM (5’-CTGCTGATGCCGCTGTTGTTCTT-3’) or IGHG (5’-GTCACCGGCTCAGGGAAGTAGC-3’).

PCR was performed using AmpliTaq Gold 360 DNA Polymerase (Applied Biosystems #43-988-23) and the following program: 7 minutes at 97°C, 40 cycles of (30 seconds at 95°C, 45 seconds at 55°C, 60 seconds at 72°C), and final extension for 10 minutes at 72°C. PCR reactions were cleaned using 1.8x CleanNGS DNA&RNA Clean-Up Magnetic Beads (Bulldog Bio #CNGS005) according to the manufacturer’s instructions. Illumina sequencing libraries were prepared from cleaned PCR reactions using the NEBNext Ultra II DNA Library Prep Kit for Illumina (NEB #E7103S) and NEBNext Multiplex Oligos for Illumina (NEB #E7600S) according to the manufacturer’s instructions. Indexed IGHM and IGHG libraries were pooled for each bat and sequenced on an Illumina MiSeq (2 x 300 bp kit).

For the captive, uninfected *A. jamaicensis* samples, IGHM and IGHG BCR repertoire sequences were obtained from Crowley *et al.* ^49^. We used BCR sequences from two individuals (B5 and B9) from experiment two of this study. Sequences and their isotype (IGHM or IGHG) were extracted from published IgBlast output files (from the “SEQUENCE” column and the “C” column, respectively) to a fasta file and used for analyses described below. Bats used in this study originated from the same colony as our RABV-infected bats.

For the wild-caught sample, RNA was extracted from a small piece of splenic tissue by homogenizing the tissue in Trizol and transferring the sample to Qiagen RNeasy columns (Qiagen #74104). 5’ RACE ready cDNA was prepared using the SMARTer RACE 5’/3’ Kit (Takara #634859) according to the manufacturer’s instructions using poly-A primers. IGH amplicons were generated via PCR with the 5’ RACE Universal Primer and custom constant region primers. Constant region primers for IGHM (5’-CCCACAGTCACMYGGCTGTCATCAGAC-3’) and IGHG (primer mix of 5’-CCAAGGACACTGTGGAGCCGGATGTG-3’ and 5’-CCAGGGACAATGTTGATCCGGATGTG-3’) were designed in Geneious (v2023.2.1) using primer3 by aligning IGHC regions from related species. Reference sequences were downloaded from IMGT ^28^ or Genbank nucleotide records and/or extracted by BLASTing ^86^ IMGT sequences to bat genomes on Genbank. Primers were designed to target the 5’ end of the CH1. Barcodes were added to amplicons from each isotype PCR by amplifying the products with primers using the same 5’ RACE Universal Primer and constant region gene-specific regions but with an added barcode and half the Illumina adapter. The second half of the adapter was added in a subsequent PCR step. Barcoded amplicons from different isotypes were pooled and sequenced on an Illumina MiSeq (2 x 300 bp kit).

### BCR-repertoire analysis

#### Pre-processing

To assess the impact of different IgBlast references on BCR-repertoire sequencing analysis, we first generated IgBlast references based on human, *E. fuscus,* or *A. jamaicensis* IGHV, IGHD, IGHJ, and IGHC gene sequences. Human sequences were obtained from IMGT ^28^, *E. fuscus* sequences from a recent annotation of the genome ^16^, and *A. jamaicensis* sequences were obtained from our genome annotation (**Table S1**). IgBlast databases for each species were generated using the *makeblastdb* command from IgBlast (v1.21.0) with arguments *-parse-seqids* and *-dbtype nucl* ^46^. For the IGHC reference database, only the CH1 sequence was considered, as the original sequencing strategy only captured the first 50-100 bp of the constant gene. An auxiliary file containing IGHJ reading frame information was also generated for each species. For each sample, IgBlast was run three times, once using human reference databases, once using *E. fuscus* reference databases, and once using *A. jamaicensis* reference databases. Sequences from the uninfected samples (from Crowley *et al.*) did not contain IGHC region sequences meaning that IGHM and IGHG isotypes could not be identified by IgBlast. Instead, isotype was determined based on sequencing primer and this information was provided with the original data. For downstream analysis, uninfected samples were separated by isotype prior to filtering. IgBlast output files were filtered, and we retained sequences meeting the following criteria: locus=IGH, productive=TRUE, complete_vdj=TRUE. Additionally, we only retained sequences with a v_call, d_call, j_call, and c_call. Finally, we filtered for unique nucleotide sequences and removed all duplicate sequences based on the sequence in the “SEQUENCE” column.

To process BCR-repertoire sequencing data from the wild-caught *A. jamaicensis*, we used AdapterRemoval (v2.3.1) ^89^ to trim adapters from all sequences and extracted reads belonging to *A. jamaicensis* by selecting sequences with the specific barcode for this individual. We used FLASH (v1.2.11) ^90^ to merge paired reads and manually removed the barcode sequences from each merged read. Pre-processed sequencing reads were assigned an IGH isotype and converted into a standardized tab-delimited database file required for downstream analysis using IgBlast (v1.21.0) and our captive *A. jamaicensis* reference database ^46^. For downstream analysis, this IgBlast output file was filtered, and we retained sequences meeting the following criteria: locus=IGH, productive=TRUE, complete_vdj=TRUE. Additionally, we only retained sequences with a v_call, d_call, j_call, and c_call. Finally, we filtered for unique nucleotide sequences and removed all duplicate sequences based on the sequence in the “SEQUENCE” column.

For both captive (RABV-infected only) and wild-caught files, isotype-specific files were generated by extracting sequences identified as IGHM or IGHG into separate files. To validate the accuracy of IGHC gene calling by IgBlast, we included the IGHE and IGHA CH1 sequences in the reference file, even though we did not amplify any IGHE or IGHA sequences in our original library. As expected, IgBlast did not identify any IGHE or IGHA sequences when files were processed with an *A. jamaicensis* reference, providing confidence in the accuracy of the program’s IGHC calls. Information about the number of sequences obtained for each sample can be found in **Table S2**.

#### Gene usage frequency

To determine IGHV and IGHJ gene usage in each isotype, we first condensed the gene-specific IGHV and IGHJ calls from the IgBlast outputs to the IGHV or IGHJ gene family. For gene calls with multiple IGHV or IGHJ genes, we only considered the first gene listed as in nearly all cases, all the genes listed belonged to the same gene family. Using the condensed IGHV and IGHJ gene calls, we counted the number of sequences for each IGHV gene family that used a particular IGHJ gene. The usage frequency of these gene pairs was then calculated for each isotype. Frequency of usage of each individual IGHV gene was also calculated and shown in **Table S3**.

#### Somatic hypermutation (SHM) frequency

SHM frequency was determined using the SHazaM module for mutation analysis, available as part of the Immcantation Portal, a collection of Python and R packages for repertoire analysis of BCR and TCR repertoires ^91^. With the isotype-specific IgBlast output files containing unique sequences, the mutation frequency across the entire IGHV region (denoted as total mutations) for replacement (R) mutations was determined using the regionDefinition argument IMGT_V_BY_SEGMENTS. For comparison of human and *A. jamaicensis* SHM frequency, human SHM values were obtained from Kitaura *et al.* (Figure 5A) ^57^ and Braddom *et al.* (Figure 1A) ^58^. From Braddom *et al.*, only total replacement SHM frequencies for naïve B cells, IGHM+ classical memory B cells, and IGHG+ classical memory B cells from U.S. donors were used for comparison with *A. jamaicensis* data. Since we were unable to differentiate between naïve B cells and IGHM class-switched B cells in the *A. jamaicensis* data, we combined the Braddom *et al.* data for SHM frequency from naïve B cells and IGHM+ classical memory B cells to compare with the IGHM *A. jamaicensis* SHM frequencies. Additional human BCR repertoire sequencing data was obtained from the SRA. IGHM and IGHG data sets from two individuals (healthy human 1: SRR7849308, SRR7849327; healthy human 2: SRR7848875, SRR7849130,) were processed as described for *A. jamaicensis* samples, with the exception that a human IgBlast reference database was used.

#### CDRH3 physicochemical properties

CDRH3 length for IGHM and IGHG sequences was calculated using the Alakazam module for amino acid physicochemical property analysis ^91^. Sequence length was determined using default arguments. The first (canonical cysteine) and last (usually tryptophan) codon were removed from the CDRH3 sequence prior to this analysis. To investigate differences in CDRH3 length related to SHM frequency in IGHM sequences, we first separated IGHM sequences based on SHM frequency into those with “low” SHM (replacement SHM < 0.01) and those with “high” SHM (replacement SHM ≥ 0.01). Then, we calculated CDRH3 length, as described above, for the low and high SHM frequency groups.

#### Cysteine-rich IGHV gene usage

Sequences from uninfected, RABV-infected, or wild-caught IGHM and IGHG BCR repertoires using artjamIGHV1.23-ctg26 were extracted from filtered, isotype-specific IgBlast files (generated as described above). To determine the frequency of these genes in the total repertoire, the number of sequences using artjamIGHV1.23-ctg26 was divided by the total number of sequences. To determine the frequency of these genes in the IGHV1-specific repertoire, the number of sequences using artjamIGHV1.23-ctg26 was divided by the total number of sequences using IGHV1 genes.

#### Repertoire diversity

Diversity of IGHM and IGHG BCR repertoires analyzed with IgBlast databases from human, *E. fuscus*, or *A. jamaicensis* references were calculated using SCOPer and the Alakazam module from the Immcantation framework ^91,92^. For each reference database (human, *E. fuscus*, and *A. jamaicensis*), we first subsampled the filtered isotype-specific IgBlast files for each experimental condition (uninfected and RABV-infected) to 1,000 sequences with replacement. Subsampled files for all samples processed with the same reference database were then combined into a single file for further analysis. Clones were assigned to sequences processed using the same IgBlast reference database with SCOPer using the argument method=”vj”. After assigning clones, clone size distribution and diversity over a range of diversity orders (q) were calculated using the *alphaDiversity* command from Alakazam. The 95% confidence interval was determined by bootstrapping (n=100). Diversity at fixed values of q was calculated using the *alphaDiversity* command with the arguments min_q=0, max_q=2, and step_q=1.

To compare the diversity of IGHM and IGHG repertoires between humans and *A. jamaicensis*, we obtained IGHM and IGHG BCR repertoire sequencing datasets from two healthy humans (healthy human 1: SRR7849308, SRR7849327; healthy human 2: SRR7848875, SRR7849130). Human datasets were pre-processed as described above. In brief, IgBlast was run using a human reference database, IgBlast output was filtered to retain unique sequences with a V, D, J, and C gene call, and files were split into isotype-specific files. We then subsampled human and *A. jamaicensis* (uninfected, RABV-infected, and wild-caught) data sets to 1,000 sequences with replacement. All subsampled data sets were combined into a single file for further analysis. Clones were assigned to sequences processed using the same IgBlast reference database with SCOPer using the argument method=”vj”. After assigning clones, clone size distribution and diversity over a range of diversity orders (q) were calculated using the *alphaDiversity* command from Alakazam. The 95% confidence interval was determined by bootstrapping (n=100). Diversity at fixed values of q was calculated using the *alphaDiversity* command with the arguments min_q=0, max_q=2, and step_q=1. This process was repeated five times, and results from all five iterations were used for Shannon diversity calculations.

### Bioinformatics

#### IGHV and IGLV gene phylogenetic analysis

To generate IGHV gene phylogenies, all functional IGHV gene translations from *A. jamaicensis* and humans (obtained from IMGT) were first aligned using the “Multiple Alignment” function (MUSCLE alignment, PPP algorithm) in Geneious (v2023.2.1). For IGLV genes, all functional IGLV gene translations from *A. jamaicensis* and all IGLV gene translations from humans (obtained from IMGT) were aligned. For IGHV and IGLV genes separately, neighbor-joining trees were constructed using the Geneious tree builder (genetic distance model=Jukes-Cantor; tree build method=neighbor-joining; no outgroup) with bootstrapping (100 replicates). The trees were then visualized and formatted using FigTree (v1.4.4) ^93^.

#### IGHC and IGLC gene alignments

Alignments of *A. jamaicensis* IGHC and IGLC genes with those from humans and *E. fuscus* were generated in Geneious (v2023.2.1) using the “Multiple Alignment” function (MUSCLE alignment, PPP algorithm).

#### IGHC gene BLAST to transcriptomics data

To validate the functionality of IGHC genes, we used transcriptomics data from a published study of *A. jamaicensis* responses to viral infection ^21^. We used BLAST ^86^ to query the coding regions of each IGHC gene (IGHM: CH1, CH2, CH3, CH4-CHS, M1; IGHG: CH1, H, CH2, CH3-CHS, M1, M2; IGHE: CH1, CH2, CH3, CH4-CHS, M1, M2; IGHA: CH1, H-CH2, CH3-CHS, M) using transcriptomic data from an uninfected *A. jamaicensis* spleen (SRA experiment set: SRX1471416) as the reference search set and optimized for highly similar sequences (megablast). Up to 100 sequences with significant alignments were generated for each query sequence.

#### Cysteine rich IGHV gene analysis

Since all functional IGHV genes require two canonical cysteines, cysteine-rich IGHV genes were considered those that contained three or more total cysteines. To identify cysteine-rich *A. jamaicensis* IGHV genes, we aligned all functional IGHV genes using the “Multiple Alignment” function (MUSCLE alignment, PPP algorithm) in Geneious (v2023.2.1) and highlighted cysteine residues.

For comparison to other species, we obtained the sequences of all functional IGHV genes for humans, rhesus monkeys, dogs, horses, cows, and pigs from IMGT ^28^. We also obtained the sequences of all functional *E. fuscus* IGHV genes from Pursell *et. al.* ^16^. IGHV sequences from each species were aligned separately using Geneious as described above for *A. jamaicensis*. For all species (including *A. jamaicensis*), IGHV genes with three or more cysteines were counted. Percentages of cysteine-rich IGHV genes were calculated by dividing the number of cysteine-rich genes by the number of functional IGHV gene sequences for each species.

#### Switch motif density

To determine the density of switch motifs preceding IGHC genes, we used Geneious (v2023.2.1) to search a region of contig ptg000026l containing the IGHJ and IGHC genes (7.27 Mb – 7.39 Mb) for the AID hotspot motif 5’-AGCT-3’ on both the forward and reverse strands ^35,36^. Raw counts were determined for non-overlapping 1.0 kb windows across the region of interest, and Z-scores for each window were calculated to obtain a normalized motif count.

## Notes

### Competing Interest Statement

The authors have declared no competing interest.

