## Supplemental Figures for "Back to basics: Immunoglobulin germline reference sequences enable investigations and reveal insights into bat-specific immunity"

### SUPPLEMENTARY FIGURES

### Figure S1

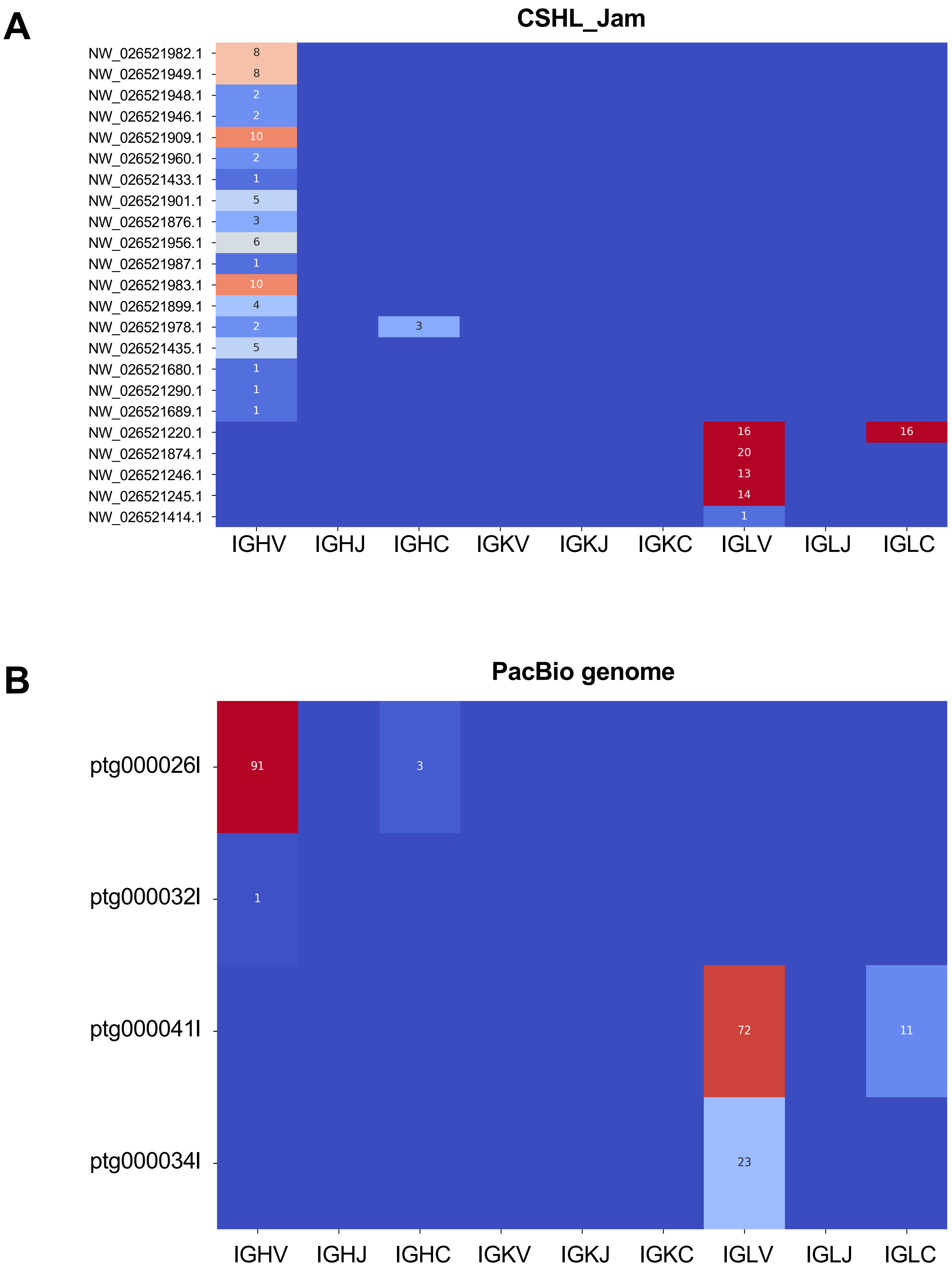

A
CSHL_Jam
NW_026521982.1
NW_026521949.1
NW_026521948.1
NW_026521946.1
NW_026521909.1
NW_026521960.1
NW_026521433.1
NW_026521901.1
NW_026521876.1
NW_026521956.1
NW_026521987.1
NW_026521983.1
NW_026521899.1
NW_026521978.1
NW_026521435.1
NW_026521680.1
NW_026521290.1
NW_026521689.1
NW_026521220.1
NW_026521874.1
NW_026521246.1
NW_026521245.1
NW_026521414.1
8
8
2
2
10
2
1
5
3
6
1
10
4
2
3
5
1
1
1
16
16
20
13
14
1
IGHV
IGHJ
IGHC
IGKV
IGKJ
IGKC
IGLV
IGLJ
IGLC
B
PacBio genome
ptg000026l
ptg000032l
ptg000041l
ptg000034l
91
3
1
72
11
23
IGHV
IGHJ
IGHC
IGKV
IGKJ
IGKC
IGLV
IGLJ
IGLC

**A** **Figure S2**

```
ArtJam-IGHM-transcript DSVSSPNLFPLISCENSLSDDSRVTVGCLAKDFLPSPVTFSWSYKNNSGISSQDIQNFPS  60
ArtJam-IGHM-M          DSVSSPNLFPLISCENSLSDDSRVTVGCLAKDFLPSPVTFSWSYKNNSGISSQDIQNFPS  60
EptFus-chr5-IGHM       ASAASPNLFPLVSCENSLSDESLVAMGCLAKDFLPDSVTFSWSFKNTTAISSQDIKNFPS  60
EptFus-chr24-IGHM      ASAAPPNLFPLVSCDNSLYDERLVAMGCLAKDFLPDSVTFSWSFKNTSAISGQDIQNFPS  60
human-IGHM-X14940      GSASAPTLFPLVSCENSPSDTSSVAVGCLAQDFLPDSITLSWKYKNNSDISST--RGFPS  58

ArtJam-IGHM-transcript VLREGKYAASSQVVLPSININQGTDEFVLCNVKHSNGDKKVQVPIQGADSGDPLSPNVNI 120
ArtJam-IGHM-M          VLREGKYAASSQVVLPSININQGTDEFVLCNVKHSNGDKKVQVPIQGADSGDPLSPNVNI 120
EptFus-chr5-IGHM       VLKEGKYEASSQVLLPTADILQGTDEFVTCKVKHSNGEKELQVALPG---GGVSPPHVNV 117
EptFus-chr24-IGHM      VLREGKYEASSQVFLPTADILQGTDNFVTCKVKHSNGEKELQVVFPG---GVVLPPHVNV 117
human-IGHM-X14940      VLRGGKYAATSQVLLPSKDVMQGTDEHVVCKVQHPNGNKEKNVPLPV---IAELPPKVSV 115

ArtJam-IGHM-transcript FIPPRDAFSGASPRKSRLICQATGFRPKKIFVSWLQDGKPLGSNFTTDKVQAEPKGPGPV 180
ArtJam-IGHM-M          FIPPRDAFSGASPRKSRLICQATGFRPKKIFVSWLQDGKPLGSNFTTDKVQAEPKGPGPV 180
EptFus-chr5-IGHM       FIPPRDSFTGPGPRTSTLICQATGFSPKKIAVSWLKEGKLLTSGFVTDKPEVERKA-RPV 176
EptFus-chr24-IGHM      FIPPRDSFTGPGPRTSTLICQATGFSPKKIAVSWLKEGKLLTSGFVTDKPEVERKA-RPM 176
human-IGHM-X14940      FVPPRDGFFG-NPRKSKLICQATGFSPRQIQVSWLREGKQVGSGVTTDQVQAEAKESGPT 174

ArtJam-IGHM-transcript TYSVSSMLMITESDWLSQSLFTCQVDHNGLSFQKNVSSVCNTGA-SNIKIFTIPPSFAST 239
ArtJam-IGHM-M          TYSVSSMLMITESDWLSQSLFTCQVDHNGLSFQKNVSSVCNTGA-SNIKIFTIPPSFAST 239
EptFus-chr5-IGHM       TYRVTSTLTITESDWLSQSVFTCQVEHQELIFQKNVSSMCGPGT-STIRVFTIPPTFASI 235
EptFus-chr24-IGHM      TYRVTSTLTITESDWLSQSVFTCQVEHQELIFQKNVSSMCGPGT-STIRVFTIPPTFASI 235
human-IGHM-X14940      TYKVTSTLTIKESDWLGQSMFTCRVDHRGLTFQQNASSMCVPDQDTAIRVFAIPPSFASI 234

ArtJam-IGHM-transcript FLTKNAKLSCLVTDLTTYDTLTISWTRQNGKDLKTHTNISEILPNATFRAVGEATVCMEE 299
ArtJam-IGHM-M          FLTKNAKLSCLVTDLTTYDTLTISWTRQNGKDLKTHTNISEILPNATFRAVGEATVCMEE 299
EptFus-chr5-IGHM       FLTKEATLSCLVTDLATYDSLRISWTRQNGDEVNTDTKVSESHPNATFSAMGKATVCVED 295
EptFus-chr24-IGHM      FLTKEATLSCLVTDLATYDSLHISWTRQNGEVMKTHTNISESHPNATFSAMGKATVCVED 295
human-IGHM-X14940      FLTKSTKLTCLVTDLTTYDSVTISWTRQNGEAVKTHTNISESHPNATFSAVGEASICEDD 294

ArtJam-IGHM-transcript WESGEEFTCTVTHADLPFPLKHTISKPKEVDKQMPSVYVLPPTREQLSLRESASVTCLVK 359
ArtJam-IGHM-M          WESGEEFTCTVTHADLPFPLKHTISKPKEVDKQMPSVYVLPPTREQLSLRESASVTCLVK 359
EptFus-chr5-IGHM       WEAGEQFTCTVTHTDLPSPLKHTIFRPKDVAKHMPSVYVQPPSREQLSLRESASVTCLVK 355
EptFus-chr24-IGHM      WEAGEQFTCTVAHTDLPSPLKHTIFRPKDVAKHMPAVYVQPPSREQLSLRESASVTCLVK 355
human-IGHM-X14940      WNSGERFTCTVTHTDLPSPLKQTISRPKGVALHRPDVYLLPPAREQLNLRESATITCLVT 354

ArtJam-IGHM-transcript GFSPPDIFVQWMQRGQPMSSDMYVTSSPLPEPQVPGRYFIHSILTVSEEDWSAGDTYTCI 419
ArtJam-IGHM-M          GFSPPDIFVQWMQRGQPMSSDMYVTSSPLPEPQVPGRYFIHSILTVSEEDWSAGDTYTCI 419
EptFus-chr5-IGHM       GFSPPDVFVQWLQRGQPVSSDMYVTSAPVPEPQAPGLYFVHSILTVSEEDWSAGETYTCV 415
EptFus-chr24-IGHM      GFSPPDVFVQWLQRGQPVSSDMYVTSAPVPEPQAPGLYFVHSILTVSEEDWSAGETYTCV 415
human-IGHM-X14940      GFSPADVFVQWMQRGQPLSPEKYVTSAPMPEPQAPGRYFAHSILTVSEEEWNTGETYTC- 413

ArtJam-IGHM-transcript VAHEALPHLVTERTVDKSTGKPTLYNVSLVISDMASTCH*                     479
ArtJam-IGHM-M          VAHEALPHLVTERTVDKSTE--------------------GEVSAEEEGFENLNTMASTF 459
EptFus-chr5-IGHM       VAHEALPYSVTERTVDKSTGKPTLYNVSLVMSDVASTCHGGEVSADEEGFENLNTMASTF 475
EptFus-chr24-IGHM      VAHEALPYSVTERTVDKSTGKPTLYNVSLVMSDVASTCHGGEVSADEEGFENLNTMASTF 475
human-IGHM-X14940      VAHEALPNRVTERTVDKSTGKPTLYNVSLVMSDTAGTCYEGEVSADEEGFENLWATASTF 473

ArtJam-IGHM-transcript
ArtJam-IGHM-M          IVLFLLSLFYSTTVTLFKVK*                                        480
EptFus-chr5-IGHM       IVLFLLSLFYSTTVTLFKVG-                                        495
EptFus-chr24-IGHM      IVLFLLSLFYSTTVTLFKVG-                                        495
human-IGHM-X14940      IVLFLLSLFYSTTVTLFKVK*                                        493
```

**B** **Figure S2**

```
ArtJam-IGHG-transcript ATTTAPKVFPLSSSCGTTSGSTVSLGCLVSGYFPEPVTVSWNSGALTRGVHTFPSILQ-S  59
ArtJam-IGHG-M          ATTTAPKVFPLSSSCGTTSGSTVSLGCLVSGYFPEPVTVSWNSGALTRGVHTFPSILQ-S  59
EptFus-chr5-IGHG       ASTTAPSVFPLSPKCGATSGSTMSLGCLVSGYFPEPVTVSWNSGALTSGVHTFPSVLR-A  59
EptFus-chr24-IGHG      ASTTAPSVFPLSPKCGATSGSMVSLGCLVSGYFPEPVTVSWNSGALTSGVHTFPSVFR-A  59
human-IGHG1-X52847     ASTKGPSVFPLAPSSKSTSGGTAALGCLVKDYFPEPVTVSWNSGALTSGVHTFPAVLQSS  60
human-IGHG2-AB006775   ASTKGPSVFPLAPCSRSTSESTAALGCLVKDYFPEPVTVSWNSGALTSGVHTFPAVLQSS  60
human-IGHG3-D78345     ASTKGPSVFPLAPCSRSTSGGTAALGCLVKDYFPEPVTVSWNSGALTSGVHTFPAVLQSS  60
human-IGHG4-AL928742   ASTKGPSVFPLAPCSRSTSESTAALGCLVKDYFPEPVTVSWNSGALTSGVHTFPAVLQSS  60
human-IGHGP-X542849    ASTKGPSVFPLVPSSRSVSEGTAALGCLVKDYFPEPVTVSWNSGALTRSVHTFPAVLQSS  60

ArtJam-IGHG-transcript GLYSLSSMVTVPASSSSSQTYTCNVDHPASSTKVEKKGESAMPRRW-------------- 105
ArtJam-IGHG-M          GLYSLSSMVTVPASSSSSQTYTCNVDHPASSTKVEKKGESAMPRRW-------------- 105
EptFus-chr5-IGHG       GLYSMSSMVTVPTTST-SQTFTCNVAHPASETKVDKIGPVTGG--P-------------- 102
EptFus-chr24-IGHG      GLYSLSSMVTVPTSSAG-QTFTCNVAHPASETKVDKIGSLTGP----------------- 101
human-IGHG1-X52847     GLYSLSSVVTVPSSSLGTQTYICNVNHKPSNTKVDKKVEPKSC--D-------------- 104
human-IGHG2-AB006775   GLYSLSSVVTVPSSNFGTQTYTCNVDHKPSNTKVDKTVERKCC----------------- 103
human-IGHG3-D78345     GLYSLSSVVTVPSSSLGTQTYTCNVNHKPSNTKVDKRVELKTPLGDTTHTCPRCPEPKSC 120
human-IGHG4-AL928742   GLYSLSSVVTVPSSSLGTKTYTCNVDHKPSNTKVDKRVESKYG----------------- 103
human-IGHGP-X542849    GLYSLSSVVTVPSSSLGTQTYTCNVDHKPSNTKVDKTVEPKTPCCD-------------- 106

ArtJam-IGHG-transcript -------------------------------SEPRCGPWQRMIEPSGGPSVFIFPPKPKD 134
ArtJam-IGHG-M          -------------------------------SEPRCGPWQRMIEPSGGPSVFIFPPKPKD 134
EptFus-chr5-IGHG       -------------------------------TPPPC-K-CPACETLGGPSVFIFPPKPKD 129
EptFus-chr24-IGHG      -------------------------------TTTPC-T-CPACETLGGPAVFIFPPKPKD 128
human-IGHG1-X52847     -------------------------------KTHTCPP-CPAPELLGGPSVFLFPPKPKD 132
human-IGHG2-AB006775   ---------------------------------VECPP-CPAP-PVAGPSVFLFPPKPKD 128
human-IGHG3-D78345     DTPPPCPRCPEPKSCDTPPPCPRCPEPKSCDTPPPCPR-CPAPELLGGPSVFLFPPKPKD 179
human-IGHG4-AL928742   ---------------------------------PPCPS-CPAPEFLGGPSVFLFPPKPKD 129
human-IGHGP-X542849    -------------------------------TTHTCPP-CATTEPLGGPSVFLFPPKPKD 134

ArtJam-IGHG-transcript ILMISRTPEVTCMVVDVAPDDLEVEFTWYVDGEKKRTAKTKAEEQQFNSTYRVVDAFSVA 194
ArtJam-IGHG-M          ILMISRTPEVTCMVVDVAPDDLEVEFTWYVDGEKKRTAKTKAEEQQFNSTYRVVDAFSVA 194
EptFus-chr5-IGHG       TLMISRTPEVTCLVVDVAPEDLDVEFTWFTDGVQGRTEKVTAQQQQFNSTYRVVNSLAIA 189
EptFus-chr24-IGHG      TLMISRTPEVTCLVVDVAPEDLDVEFTWYMDGAHVRTEKVTAQQQQFNSTYRVVNSLAIM 188
human-IGHG1-X52847     TLMISRTPEVTCVVVDVSHEDPEVKFNWYVDGVEVHNAKTKPREEQYNSTYRVVSVLTVL 192
human-IGHG2-AB006775   TLMISRTPEVTCVVVDVSHEDPEVQFNWYVDGVEVHNAKTKPREEQFNSTFRVVSVLTVV 188
human-IGHG3-D78345     TLMISRTPEVTCVVVDVSHEDPEVQFKWYVDGVEVHNAKTKPREEQYNSTFRVVSVLTVL 239
human-IGHG4-AL928742   TLMISRTPEVTCVVVDVSQEDPEVQFNWYVDGVEVHNAKTKPREEQFNSTYRVVSVLTVL 189
human-IGHGP-X542849    TLMISRTPEVTCVVVDVSHEDPEVKFNWYVDGVEVHNAKTKPWEEQYNSTYHVVSVLTVV 194

ArtJam-IGHG-transcript HQDWLNGKKFKCKVNSGSLPSPIEKTISKTPGQAQEPQVYVLAPHMDELSKSTVSVTCLV 254
ArtJam-IGHG-M          HQDWLNGKKFKCKVNSGSLPSPIEKTISKTPGQAQEPQVYVLAPHMDELSKSTVSVTCLV 254
EptFus-chr5-IGHG       HQDWLNGKEFKCKVNNKAIPAPIERTISKARGQAREPQVYVLGPHADEMAKDKVSVTCLI 249
EptFus-chr24-IGHG      HQDWLKGKEFKCKVNNNAIPAPVERTISKARGQLREPQVYVLGPHADEMAKDKVSVTCLV 248
human-IGHG1-X52847     HQDWLNGKEYKCKVSNKALPAPIEKTISKAKGQPREPQVYTLPPSRDELTKNQVSLTCLV 252
human-IGHG2-AB006775   HQDWLNGKEYKCKVSNKGLPAPIEKTISKTKGQPREPQVYTLPPSREEMTKNQVSLTCLV 248
human-IGHG3-D78345     HQDWLNGKEYKCKVSNKALPAPIEKTISKTKGQPREPQVYTLPPSREEMTKNQVSLTCLV 299
human-IGHG4-AL928742   HQDWLNGKEYKCKVSNKGLPSSIEKTISKAKGQPREPQVYTLPPSQEEMTKNQVSLTCLV 249
human-IGHGP-X542849    HQNWLNGREYKCKVSNKGLPAPIEKTISKTKGQPREPQVYTLPPSQ-KMTKNQVTLTCLV 253
```

#### Figure S2

```
ArtJam-IGHG-transcript KDFYPAEINVEWQNNGKPEPETKYTTTPPMKDKDGSYFLYSKLSVEKARWMQGQTFTCAV   314
ArtJam-IGHG-M          KDFYPAEINVEWQNNGKPEPETKYTTTPPMKDKDGSYFLYSKLSVEKARWMQGQTFTCAV   314
EptFus-chr5-IGHG       KDFFPPDISVEWQSSGRPEPETKYSSTPPQKDQEGAFFLYSKLSVDKARWQRGDPFTCEV   309
EptFus-chr24-IGHG      KDFFPPDISVEWQSNGRPEPETKYSSTPPQKDQEGAFFLYSKLSVDKARWQRGDPFMCEV   308
human-IGHG1-X52847     KGFYPSDIAVEWESNGQP--ENNYKTTPPVLDSDGSFFLYSKLTVDKSRWQQGNVFSCSV   310
human-IGHG2-AB006775   KGFYPSDIAVEWESNGQP--ENNYKTTPPMLDSDGSFFLYSKLTVDKSRWQQGNVFSCSV   306
human-IGHG3-D78345     KGFYPSDIAVEWESSGQP--ENNYNTTPPMLDSDGSFFLYSKLTVDKSRWQQGNIFSCSV   357
human-IGHG4-AL928742   KGFYPSDIAVEWESNGQP--ENNYKTTPPVLDSDGSFFLYSRLTVDKSRWQEGNVFSCSV   307
human-IGHGP-X542849    KGFYPSDITVEWESNGQP--ENNYKTTPPMLDSNGSFFLYSKLTVDKSRWQQGNVFSCSV   311

ArtJam-IGHG-transcript MHEALHNHYTQKSISQTPGK*                                          373
ArtJam-IGHG-M          MHEALHNHYTQKSISQTPGM---LLDESCAEAQDGELDGLWTTISIFITLFLLSVCYSAT   371
EptFus-chr5-IGHG       MHEALHNHYTQKTVSRSPGKTELLLDESCAEAQDGELDGLWTTISIFITLFLLSVCYSAT   369
EptFus-chr24-IGHG      MHEALHNHYAQKSVSRSPGK-ELLLDESCAEAQDGELDGLWTTISIFITLFLLSVCYSAT   367
human-IGHG1-X52847     MHEALHNHYTQKSLSLSPGK-ELQLEESCAEAQDGELDGLWTTITIFITLFLLSVCYSAT   369
human-IGHG2-AB006775   MHEALHNHYTQKSLSLSPGK-ELQLEESCAEAQDGELDGLWTTITIFITLFLLSVCYSAT   365
human-IGHG3-D78345     MHEALHNRFTQKSLSLSPGK-ELQLEESCAEAQDGELDGLWTTITIFITLFLLSVCYSAT   416
human-IGHG4-AL928742   MHEALHNHYTQKSLSLSLGK-ELQLEESCAEAQDGELDGLWTTITIFITLFLLSVCYSAT   366
human-IGHGP-X542849    MHEGLHNHYTQKSLSLSPGK-ELQLEESCAEAQDGELDGLWTTITIFITLFLLSVCYSAT   370

ArtJam-IGHG-transcript
ArtJam-IGHG-M          VTLFKVKWIFSSVVELKRTIVPDYRNMIGQGA*                              404
EptFus-chr5-IGHG       VTLFKVKWIFSSVADLKRTIVPDYRNMIGQGA*                              402
EptFus-chr24-IGHG      VTLFKVKWIFSSVADLKRTIVPDYRNMIGQGA*                              400
human-IGHG1-X52847     VTFFKVKWIFSSVVDLKQTIIPDYRNMIGQGA*                              401
human-IGHG2-AB006775   ITFFKVKWIFSSVVDLKQTIVPDYRNMIRQGA*                              397
human-IGHG3-D78345     VTFFKVKWIFSSVVDLKQTIIPDYRNMIGQGA*                              448
human-IGHG4-AL928742   VTFFKVKWIFSSVVDLKQTIVPDYRNMIRQGA*                              398
human-IGHGP-X542849    VTFFKVKWIFSSVVDLKQTIVPDYRNMIGQGA*                              402
```

**C** **Figure S2**

```
ArtJam-IGHE-transcript VSRQHPSIFPLVPCHSSISMGDTSVTLGCLIKDYIPEPVNVTWDAGSLEHSVMTLPGTLD  60
ArtJam-IGHE-M          VSRQHPSIFPLVPCHSSISMGDTSVTLGCLIKDYIPEPVNVTWDAGSLEHSVMTLPGTLD  60
EptFus-chr5-IGHE       ATVQGPSVFPLAPCSKGPAGGAASVTLGCLVRDYFPEPVTVAWDAGSLTTSVVTFPAAFN  60
EptFus-chr24-IGHE      ASMQGPSVFPLAPCSKSPAGGAASVTLGCLVRDYFPEPVTVTWDAGSLTTSVVTLPATFS  60
human-IGHE-J00222      ASTQSPSVFPLTRCCKNIPSNATSVTLGCLATGYFPEPVMVTCDTGSLNGTTMTLPATTL  60

ArtJam-IGHE-transcript STTGLYTTISQTTTSGEWATQKFICSVEHDGAAVEK-TISVRKVCATNVTQPTVKLYHSS  119
ArtJam-IGHE-M          STTGLYTTISQTTTSGEWATQKFICSVEHDGAAVEK-TISVRKVCATNVTQPTVKLYHSS  119
EptFus-chr5-IGHE       --SGLYTTSSQVTASGEWAKQRFTCSVVHPAGSTTV-NRTIDPACATNVTLPTVGIYHSS  117
EptFus-chr24-IGHE      --SGLYTTSSQVTASGEWAKQRFTCSVAHPAGSTTV--NKTIDACATNVTLPTVGIYHSS  116
human-IGHE-J00222      TLSGHYATISLLTVSGAWAKQMFTCRVAHTPSSTDWVDNKTFSVCSRDFTPPTVKILQSS  120

ArtJam-IGHE-transcript CNPSGNTQATIQLLCLISDFTPGDIKVTWLVDGHMDKSMFPYTSTPKQEGNLSSIFSQLN  179
ArtJam-IGHE-M          CNPSGNTQATIQLLCLISDFTPGDIKVTWLVDGHMDKSMFPYTSTPKQEGNLSSIFSQLN  179
EptFus-chr5-IGHE       CDPSGNTQATIQLLCLIPGFTPGDLEVTWLVDGQTE-NLFPITGPASWEGALASTFSQLN  176
EptFus-chr24-IGHE      CDPSGNTQATIQLLCLIPGFTPGDLEVTWLVDGQTE-NLFPITGPASWEGALASTFSQLN  175
human-IGHE-J00222      CDGGGHFPPTIQLLCLVSGYTPGTINITWLEDGQVM-DVDLSTASTTQEGELASTQSELT  179

ArtJam-IGHE-transcript ITQGEWVSQKTYTCQVYYQGCKFEKHARNCP-ESEPRGVSVYLIPPSPLDLYLHKSPKIT  238
ArtJam-IGHE-M          ITQGEWVSQKTYTCQVYYQGCKFEKHARNCP-ESEPRGVSVYLIPPSPLDLYLHKSPKIT  238
EptFus-chr5-IGHE       ISQGEWLSQRTYACQASLHGCTVKAEARACPPEAEPRGVSVYVIPPSPLDLYVHKSPKVT  236
EptFus-chr24-IGHE      ISQGEWLSQRTYACQASLHGCTVKAEARACPPESEPRGVSVYVIPPSPLDLYVHKSPKVA  235
human-IGHE-J00222      LSQKHWLSDRTYTCQVTYQGHTFEDSTKKCA-DSNPRGVSAYLSRPSPFDLFIRKSPTIT  238

ArtJam-IGHE-transcript CLVVDLASID-SVTLQWFRESRGLVKDTMWNSKLQFNMTYTVTSTLPVEANDWIEGETYE  297
ArtJam-IGHE-M          CLVVDLASID-SVTLQWFRESRGLVKDTMWNSKLQFNMTYTVTSTLPVEANDWIEGETYE  297
EptFus-chr5-IGHE       CLVVDLASLEGMS-LQWSREGGDLTNEATQTNKRHFNMTYSVTSTLPVDAGDWIEGETYK  295
EptFus-chr24-IGHE      CLAVDLASLEGMS-LQWSREGGGLLTEATQSSKRHFNMTYTVTSTLPVDAGDWIEGETYE  294
human-IGHE-J00222      CLVVDLAPSKGTVNLTWSRASGKPVNHSTRKEEKQRNGTLTVTSTLPVGTRDWIEGETYQ  298

ArtJam-IGHE-transcript CRLTHPHLPREIVRTISKGNGKRVVPEVYVFLPPEEEQGTKDTLTLTCLIQNFFPADISV  357
ArtJam-IGHE-M          CRLTHPHLPREIVRTISKGNGKRVVPEVYVFLPPEEEQGTKDTLTLTCLIQNFFPADISV  357
EptFus-chr5-IGHE       CKLTHPDLPQDIVRTIAKAPGKRAAPEVFVFPPPKEE-GAQDTLTLTCLIQGFFPADISV  354
EptFus-chr24-IGHE      CRLSHPDLPQDIVRTIAKAPGKRAAPEVFLFPPPK-EQGAEDTLTLTCLIQGFFPADVSV  353
human-IGHE-J00222      CRVTHPHLPRALMRSTTKTSGPRAAPEVYAFATPE-WPGSRDKRTLACLIQNFMPEDISV  357

ArtJam-IGHE-transcript QWLCNNSLIPTNQQATTWPLKVNSSSRTFFIFSRLEVSRADWEKRSQFTCQVVHEALSGS  417
ArtJam-IGHE-M          QWLCNNSLIPTNQQATTWPLKVNSSSRTFFIFSRLEVSRADWEKRSQFTCQVVHEALSGS  417
EptFus-chr5-IGHE       QWLRDNALLPSHQYTTTRPLRDRGASPAFFVFSRLEVKGADW-KRSNFTCRVTHEALPNS  413
EptFus-chr24-IGHE      QWLQDDALIPSNQHATTRPLRDPGTSPAFFLFSRLEVSRADWEKRSNFTCRVTHEALPHS  376
human-IGHE-J00222      QWLHNEVQLPDARHSTTQPRKTKGS--GFFVFSRLEVTRAEWEQKDEFICRAVHEAASPS  415

ArtJam-IGHE-transcript RTLEKSVSRGPGN*                                               477
ArtJam-IGHE-M          RTLEKSVSRGPEVD--LQDLCAGVAESEELDGIWTSLFIFVVLFLLSVTYGASVTLFK   475
EptFus-chr5-IGHE       RTHDQSVSKGPGNELDLQDLCVGAAEAEELDGLWRSLLVFIALFLLSVTYGASVTLFKAK  473
EptFus-chr24-IGHE      RTIKQSVSKDSGNELDLQDLCVGAAEGEELDGLWRSLFVFIALFLLSVTYGASVTLFKAK  420
human-IGHE-J00222      QTVQRAVSVNPGKE--L-DVCVEEAEGEA-PWTWTGLCIFAALFLLSVSYSAALTLLMVQ  471

ArtJam-IGHE-transcript
ArtJam-IGHE-M
EptFus-chr5-IGHE
EptFus-chr24-IGHE      WVLDAVLQGPPQATHDYENIEDMAQ                                    445
human-IGHE-J00222      RFLSATRQGRPQTSLDYTNVLQPHA                                    496
```

# D

### Figure S2

```
ArtJam-IGHA-transcript VDPARPSSFPLSLQSTDASGHVVIGCLVQGFFPPGSVNVTWDHNGEGTSIRNFPPVQAAG  60
ArtJam-IGHA-M          VDPARPSSFPLSLQSTDASGHVVIGCLVQGFFPPGSVNVTWDHNGEGTSIRNFPPVQAAG  60
EptFus-chr5-IGHA       NNPTSPTLFPLTFRSTKPSDPVIIGCLVQDFLPSKPVDVTWDSSGSGTSISNVLPMFTSR  60
eptfus-chr24-IGHA      ANPSSPTLFPLSLRSTDPAGHVVIGCLVQDFFPPEPMKVTWGSSGPGTSIRNFLPVLNSR  60
human-IGHA1-BK063801   ASPTSPKVFPLSLCSTQPDGNVVIACLVQGFFPQEPLSVTWSESGQGVTARNFPPSQDAS  60
human-IGHA2-BK063800   ASPTSPKVFPLSLDSTPQDGNVVVACLVQGFFPQEPLSVTWSESGQNVTARNFPPSQDAS  60

ArtJam-IGHA-transcript -GLY-TMSSQLTLPADQCPKKGTQKCHVKYNSSPSQAVNVPCS----TE----------- 103
ArtJam-IGHA-M          -GLY-TMSSQLTLPADQCPKKGTQKCHVKYNSSPSQAVNVPCS----TE----------- 103
EptFus-chr5-IGHA       -ARGTEVSASLSVPPTPMP--------------------------TS-----------  80
eptfus-chr24-IGHA      -GLY-TMVSQLTLPADQCPASATQKCHVQHNSSPSQTKDVPCKGQSVTP----------- 107
human-IGHA1-BK063801   GDLY-TTSSQLTLPATQCLAGKSVTCHVKHYTNPSQDVTVPCP----VPSTPPTPSPSTP 115
human-IGHA2-BK063800   GDLY-TTSSQLTLPATQCPDGKSVTCHVKHYTNPSQDVTVPCP----VP----------- 104

ArtJam-IGHA-transcript -KCKGSCCQPHLTLHPPTLEDLLLGFNANITCTLSGLQDPKDATFTWKPTGGKEAIQGTT 162
ArtJam-IGHA-M          -KCKGSCCQPHLTLHPPTLEDLLLGFNANITCTLSGLQDPKDATFTWKPTGGKEAIQGTT 162
EptFus-chr5-IGHA       -RPTCSCCQPRLTLHPPALEDLLLGSDANLTCTLSGLREPQGASFTWQPTGGKKAIQQDP 139
eptfus-chr24-IGHA      -KPTSSCCQPRLTLHPPALEDLLLGSNANLTCTLSGLREPQGASFTWQPTGGKEAIQEAP 166
human-IGHA1-BK063801   PTPSPSCCHPRLSLHRPALEDLLLGSEANLTCTLTGLRDASGVTFTWTPSSGKSAVQGPP 175
human-IGHA2-BK063800   --PPPPCCHPRLSLHRPALEDLLLGSEANLTCTLTGLRDASGATFTWTPSSGKSAVQGPP 162

ArtJam-IGHA-transcript ERDSFGCYSVTSILPGCAELWNRGDTFSCTVTHPESKSPLTATITKTSGSLLPPQVHLLP 222
ArtJam-IGHA-M          ERDSFGCYSVTSILPGCAELWNRGDTFSCTVTHPESKSPLTATITKTSGSLLPPQVHLLP 222
EptFus-chr5-IGHA       QSDACGCYSVSSVLPGCADPWNRGQRFSCTATHPEATSALT-------GSTFPPQVHLLP 192
eptfus-chr24-IGHA      QRDACGCYSVASVLPGCAEPWNRGQRFSCTAAHPEAASPLTATITRASGSTFPPQVHLLP 226
human-IGHA1-BK063801   ERDLCGCYSVSSVLPGCAEPWNHGKTFTCTAAYPESKTPLTATLSK-SGNTFRPEVHLLP 234
human-IGHA2-BK063800   ERDLCGCYSVSSVLPGCAQPWNHGETFTCTAAHPELKTPLTANITK-SGNTFRPEVHLLP 221

ArtJam-IGHA-transcript PPSEELALNEMVTLTCLVRGFSPRDVLVLWRHGDQELSREKYLTWSPLKEDVQG-TMFAV 281
ArtJam-IGHA-M          PPSEELALNEMVTLTCLVRGFSPRDVLVLWRHGDQELSREKYLTWSPLKEDVQG-TMFAV 281
EptFus-chr5-IGHA       PPSEELALNELVTLTCVVRGFYPKDVLVRWLHESQELPREKYQTWSPLPEAGQGGATFAT 252
eptfus-chr24-IGHA      PPSEELALNELVTLTCVVRGFYPKDVLVRWLHESQELPREKYQTWSPLPEAGQGGATFAT 286
human-IGHA1-BK063801   PPSEELALNELVTLTCLARGFSPKDVLVRWLQGSQELPREKYLTWASRQEPSQGTTTFAV 294
human-IGHA2-BK063800   PPSEELALNELVTLTCLARGFSPKDVLVRWLQGSQELPREKYLTWASRQEPSQGTTTFAV 281

ArtJam-IGHA-transcript TSVLRVESEHWKNGDNFSCMVGHEALPRNFTQKTIDRLAGKPTHVNVSVILSEVDGTCY* 340
ArtJam-IGHA-M          TSVLRVESEHWKNGDNFSCMVGHEALPRNFTQKTIDRLAGSQ------------------ 323
EptFus-chr5-IGHA       TSVLRVDTEAWKSGDNYSCMVGHEALPLSFTQKTINRLAGKPTHVNVSVVMSEVDGVCG- 311
eptfus-chr24-IGHA      TSVLRVDTEAWKNGDNYSCMVGHEALPLSFTQKTINRLAGKPTHVNVSVVMSEVDGICY* 345
human-IGHA1-BK063801   TSILRVAAEDWKKGDTFSCMVGHEALPLAFTQKTIDRLAGKPTHVNVSVVMAEVDGTCYG 354
human-IGHA2-BK063800   TSILRVAAEDWKKGDTFSCMVGHEALPLAFTQKTIDRLAGKPTHVNVSVVMAEVDGTCYG 341

ArtJam-IGHA-transcript
ArtJam-IGHA-M          --CLADYQGPLSWVVMDLPQDNLEEDTPGASLWPTTVTLLTLFLLSLFYSMALTVTSIRG 381
EptFus-chr5-IGHA       SRCLAGCQELLPCVLIDLPQENREEDAPGASLWPTTVTLLTLFLLSLFYSTALTVTSVRG 371
eptfus-chr24-IGHA
human-IGHA1-BK063801   SCSVADWQMPPPYVVLDLPQETLEEETPGANLWPTTITFLTLFLLSLFYSTALTVTSVRG 414
human-IGHA2-BK063800   SCCVADWQMPPPYVVLDLPQETLEEETPGANLWPTTITFLTLFLLSLFYSTALTVTSVRG 401

ArtJam-IGHA-transcript
ArtJam-IGHA-M          PPSSREGPQY*                                                  392
EptFus-chr5-IGHA       PPSSREGPPF-                                                  381
eptfus-chr24-IGHA
human-IGHA1-BK063801   PSGNREGPQY*                                                  424
human-IGHA2-BK063800   PSGKREGPQY*                                                  411
```

### Figure S3

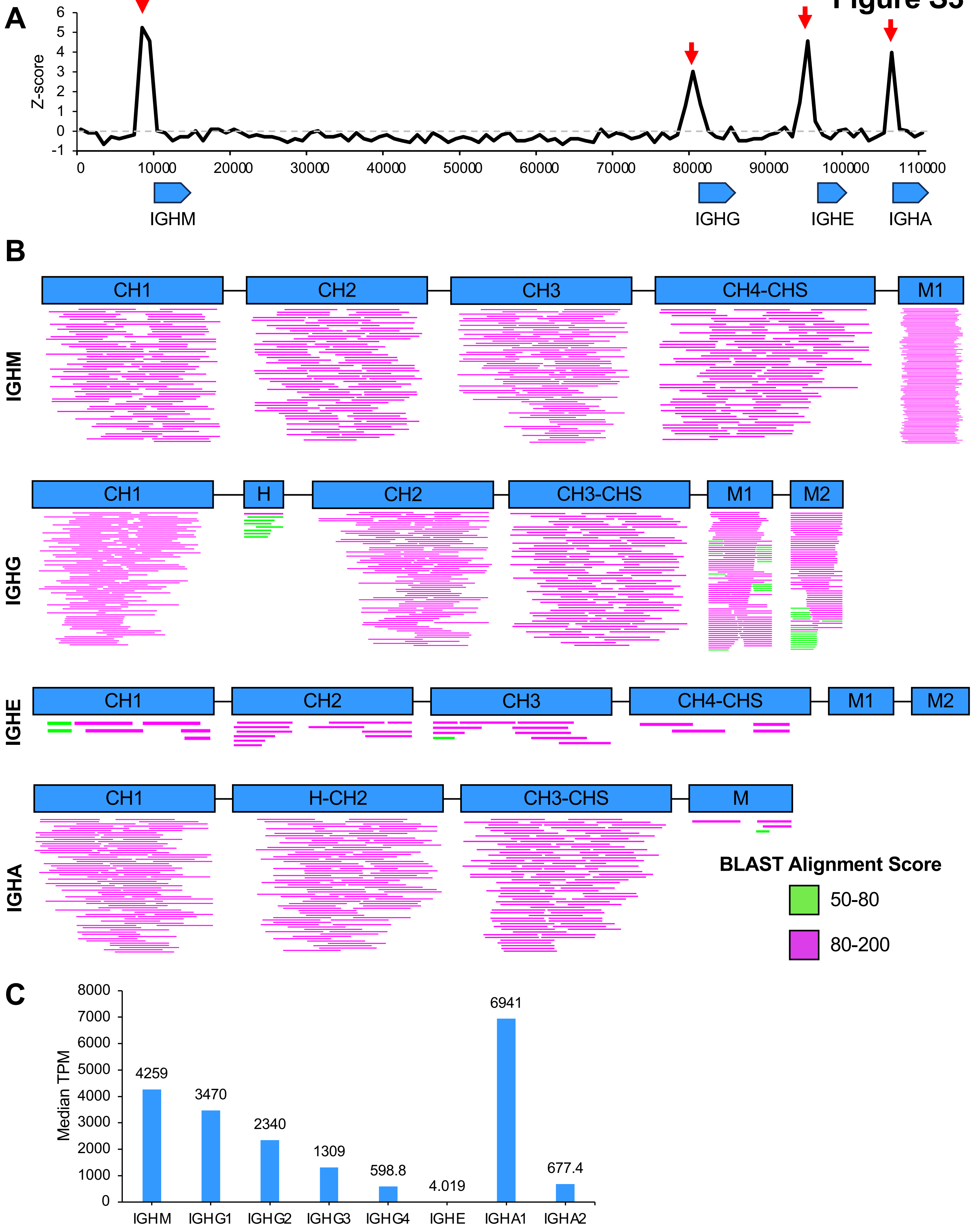
A
Z-score
0
10000
20000
30000
40000
50000
60000
70000
80000
90000
100000
110000
IGHM
IGHG
IGHE
IGHA
B
IGHM
CH1
CH2
CH3
CH4-CHS
M1
IGHG
CH1
H
CH2
CH3-CHS
M1
M2
IGHE
CH1
CH2
CH3
CH4-CHS
M1
M2
IGHA
CH1
H-CH2
CH3-CHS
M
BLAST Alignment Score
50-80
80-200
C
Median TPM
4259
3470
2340
1309
598.8
4.019
6941
677.4
IGHM
IGHG1
IGHG2
IGHG3
IGHG4
IGHE
IGHA1
IGHA2

### Figure S4

| | | |
|---|---|---|
| ArtJam-IGLC1 | GQPKSAPSVTLFPPSSEELSNNKATLVCMINGFYPEDVTVTWKENGSPVFQGVETTKPSK | 60 |
| ArtJam-IGLC2 | GQPKSAPSVTLFPPSTEELSNNKATLVCLVSDFYPGAVTVAWKENGSPVSQGVETTKPSK | 60 |
| ArtJam-IGLC3 | GQPKSAPSVTLFPPSSEELSNNKATLVCLISDFYPGAVTVAWKENGSPVSQGVETTKPSK | 60 |
| ArtJam-IGLC4 | GQPKSAPSVTLFPPSSEELSTNKATLVCLVSDFYPGAVTVAWKENGSPVSQGVETTKPSK | 60 |
| ArtJam-IGLC5 | GQPKSAPSVTLFPPSSEELSNNKATLVCLVSDFYPGAVTVAWKENGSPVSQGVETTKPSK | 60 |
| ArtJam-IGLC6 | GQPKSAPSVTLFPPSSEELSNNKATLVCLVSDFYPGAVTVAWKENGSPVSQGVETTKPSK | 60 |
| ArtJam-IGLC7 | GQPKSAPSVTLFPPSSEELSTNKATLVCLVSDFYPGAVTVAWKENGSPVSQGVETTKPSK | 60 |
| ArtJam-IGLC8 | GQPKSAPSVTLFPPSSEELSTNKATLVCLVSDFYPGAVTVAWKENGSPVSQGVETTKPSK | 60 |
| ArtJam-IGLC9 | GQPKSAPSVTLFPPSSEELSTNKATLVCLVSDFYPGAVTVAWKENGSPVSQGVETTKPSK | 60 |
| ArtJam-IGLC10 | GQPKSAPSVTLFPPSSEELSNNKATLVCLVSDFYPGAVTVAWKENGSPVSQGVETTKPSK | 60 |
| EptFus-IGLC1 | GQPKSAPSVSLFPPSAEELSKNKATLVCLMSDFYPGSVTVAWKANGTPVTQGVETTKPSK | 60 |
| EptFus-IGLC2 | GQPKSAPSVSLFPPSAEELSKNKATLVCLMSDFYPGSVTVAWKANGTPVTQGVETTKPSK | 60 |
| EptFus-IGLC3 | GQPKSEPSVSLFPPSAEELSKNKATLVCLMSDFYPGSVTVAWKANGTPVTQGVETTKPSK | 60 |
| EptFus-IGLC4 | GQPKSEPSVSLFPPSAEELSKNKATLVCLMSDFYPGSVTVAWKANGTPVTQGVETTKPSK | 60 |
| EptFus-IGLC5 | GQPKSAPSVSLFPPSAEELSKNKATLVCLMSDFYPGSVTVAWKANGTPVTQGVETTKPSK | 60 |
| EptFus-IGLC6 | GQPKSAPSVSLFPPSAEELSKNKATLVCLMSDFYPGSVTVAWKANGTPVTQGVETTKPSK | 60 |
| EptFus-IGLC7 | GQPKSEPSVSLFPPSAEELSKNKATLVCLMSDFYPGSVTVAWKANGTPVTQGVETTKPSK | 60 |
| EptFus-IGLC8 | GQPKSEPSVSLFPPSTEELSKNKATLVCLMSDFYPGSVTVAWKANGTPVTQGVETTKPSK | 60 |
| EptFus-IGLC9 | GQPKSAPSVSLFPPSAEELSKNKATLVCLMSDFYPGSVTVAWKANGTPVTQGVETTKPSK | 60 |
| EptFus-IGLC10 | GQPKSAPSVSLFPPSAEELSKNKATLVCLMSDFYPGSVTVAWKANGTPVTQGVETTKPSK | 60 |
| EptFus-IGLC11 | GQPKSAPSVSLFPPSTEELSKNKATLVCLMSDFYPGSVTVAWKANGTPVTQGVETTKPSK | 60 |
| EptFus-IGLC12 | GQPKSAPSVSLFPPSAEELSKNKATLVCLMSDFYPGSVTVAWKANGTPVTQGVETTKPSK | 60 |
| human-IGLC1-X51755 | GQPKANPTVTLFPPSSEELQANKATLVCLISDFYPGAVTVAWKADGSPVKAGVETTKPSK | 60 |
| human-IGLC2-J00253 | GQPKAAPSVTLFPPSSEELQANKATLVCLISDFYPGAVTVAWKADSSPVKAGVETTTPSK | 60 |
| human-IGLC3-K01326 | GQPKAAPSVTLFPPSSEELQANKATLVCLISDFYPGPVTVAWKADSSPVKAGVETTTPSK | 60 |
| human-IGLC6-J03011 | GQPKAAPSVTLFPPSSEELQANKATLVCLISDFYPGAVKVAWKADGSPVNTGVETTTPSK | 60 |
| human-IGLC7-X51755 | GQPKAAPSVTLFPPSSEELQANKATLVCLVSDFYPGAVTVAWKADGSPVKVGVETTKPSK | 60 |

| | | |
|---|---|---|
| ArtJam-IGLC1 | QSNNKYAASSYLSMSASKWKSASLFSCQVTHDGSTMEKTVVPSECS | 106 |
| ArtJam-IGLC2 | QSNNKYAASSYLSMSASKWKSASQFSCHVTHDGTTVDKTVVPSECS | 106 |
| ArtJam-IGLC3 | QSNNKYAASSYLSMSASKWKSASQFSCHVTHDGATVDKTVVPSECS | 106 |
| ArtJam-IGLC4 | QSNNKYAASSYLSMSASKWKSASQFSCHVTHDGTTVDKTVVPSECS | 106 |
| ArtJam-IGLC5 | QSNNKYAASSYLSMSASKWKSASQFSCHVTHDGTTVDKTVVPSECS | 106 |
| ArtJam-IGLC6 | QSNNKYAASSYLSVSASKWKSASQFSCHVTHDGTTVDKTVVPSECS | 106 |
| ArtJam-IGLC7 | QSNNKYAASSYLSMSASKWKSASQFSCHVTHDGTTVDKTVVPSECS | 106 |
| ArtJam-IGLC8 | QSNNKYAASSYLSMSASKWKSASQFSCHVTHDGTTVDKTVVPSECS | 106 |
| ArtJam-IGLC9 | QSNNKYAASSYLSMSASKWKSASQFSCHVTHDGTTVDKTVVPSECS | 106 |
| ArtJam-IGLC10 | QSNNKYAASSYLSMSASKWKSASQFSCHVTHDGTTVDKTVVPSECS | 106 |
| EptFus-IGLC1 | QSNNKYAASSYLSVSSQDWKSASAYSCQVTHDGKTVEKTVAPSECS | 106 |
| EptFus-IGLC2 | QSNNKYAASSYLSVSSQDWKSASAYSCQVTHDGKTVEKTVAPSECS | 106 |
| EptFus-IGLC3 | QSNNKYAASSYLSVSSQDWKSASAYSCQVTHDGKTVEKTVAPSECS | 106 |
| EptFus-IGLC4 | QSNNKYAASSYLSVSSQDWKSASAYSCQVTHDGKTVEKTVAPSECS | 106 |
| EptFus-IGLC5 | QSNNKYAASSYLSVSSQDWKSASAYSCQVTHDGKTVEKTVAPSECS | 106 |
| EptFus-IGLC6 | QSNNKYAASSYLSVSSQDWKSASAYSCEVTHDGKTVEKTVAPSECS | 106 |
| EptFus-IGLC7 | QSNNKYAASSYLSVSSQDWKSASAYSCQVTHDGKTVEKTVAPSECS | 106 |
| EptFus-IGLC8 | QSNNKYAASSYLSVSSQDWKSASAYSCQVTHDGKTVEKTVAPSECS | 106 |
| EptFus-IGLC9 | QSNNKYAASSYLSVSSQDWKSASAYSCQVTHDGKTVEKTVAPSECS | 106 |
| EptFus-IGLC10 | QSNNKYAASSYLSVSSQDWKSASAYSCQVTHDGKTVEKTVAPSECS | 106 |
| EptFus-IGLC11 | QSNNKYAASSYLSVSSQDWKSASAYSCEVTHDGKTVEKTVAPSECS | 106 |
| EptFus-IGLC12 | QSNNKYAASSYLSVSSQDWKSASAYSCEVTHDGKTVEKTVAPSECS | 106 |
| human-IGLC1-X51755 | QSNNKYAASSYLSLTPEQWKSHRSYSCQVTHEGSTVEKTVAPTECS | 106 |
| human-IGLC2-J00253 | QSNNKYAASSYLSLTPEQWKSHRSYSCQVTHEGSTVEKTVAPTECS | 106 |
| human-IGLC3-K01326 | QSNNKYAASSYLSLTPEQWKSHKSYSCQVTHEGSTVEKTVAPTECS | 106 |
| human-IGLC6-J03011 | QSNNKYAASSYLSLTPEQWKSHRSYSCQVTHEGSTVEKTVAPAECS | 106 |
| human-IGLC7-X51755 | QSNNKYAASSYLSLTPEQWKSHRSYSCRVTHEGSTVEKTVAPAECS | 106 |

### Figure S5

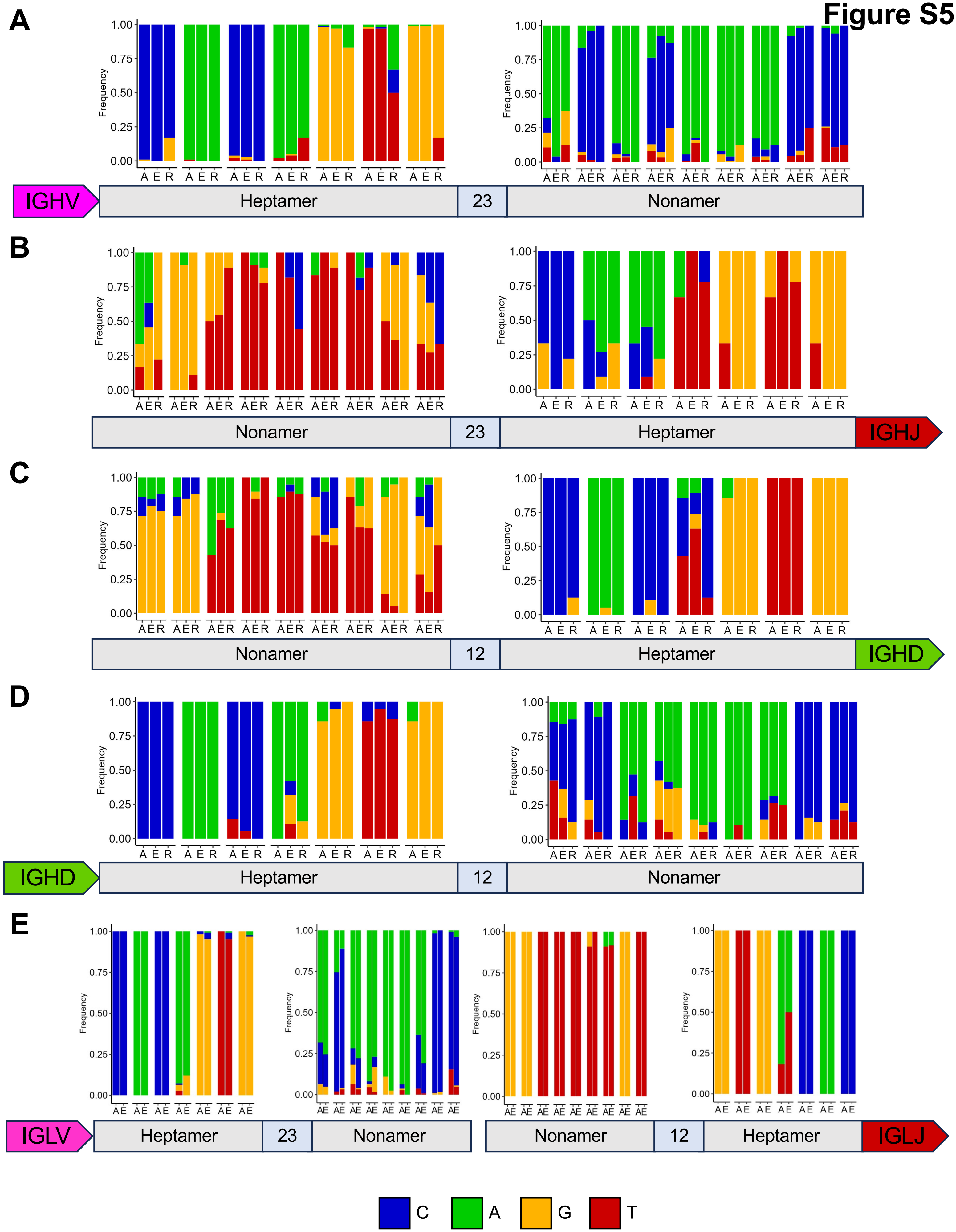
A
B
C
D
E
Frequency
1.00
0.75
0.50
0.25
0.00
A E R
AE
IGHV
Heptamer
23
Nonamer
IGHJ
12
IGHD
IGLV
IGLJ
C
A
G
T

### Figure S6

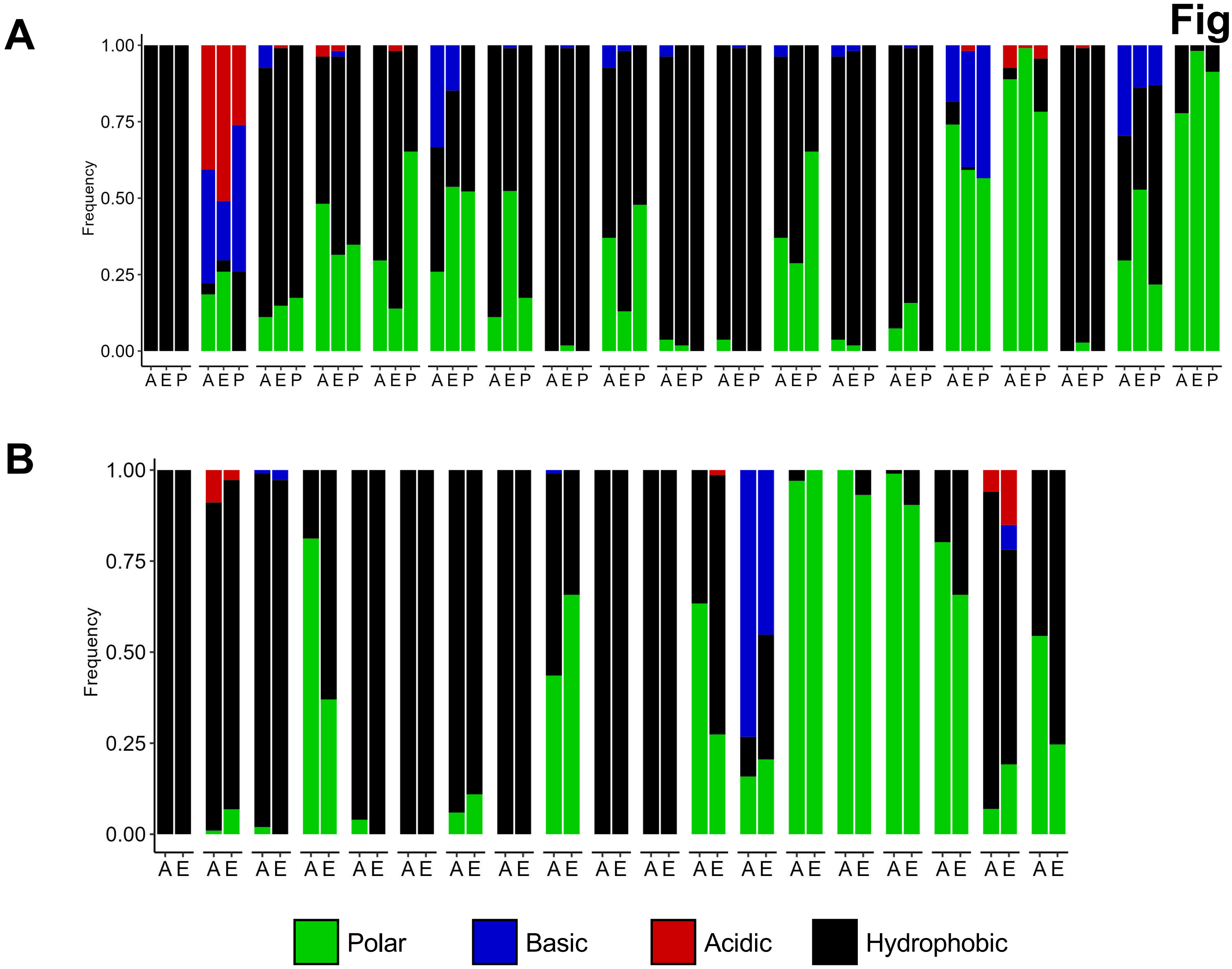
A
B
Frequency
1.00
0.75
0.50
0.25
0.00
A E P
A E
Polar
Basic
Acidic
Hydrophobic

### Figure S7

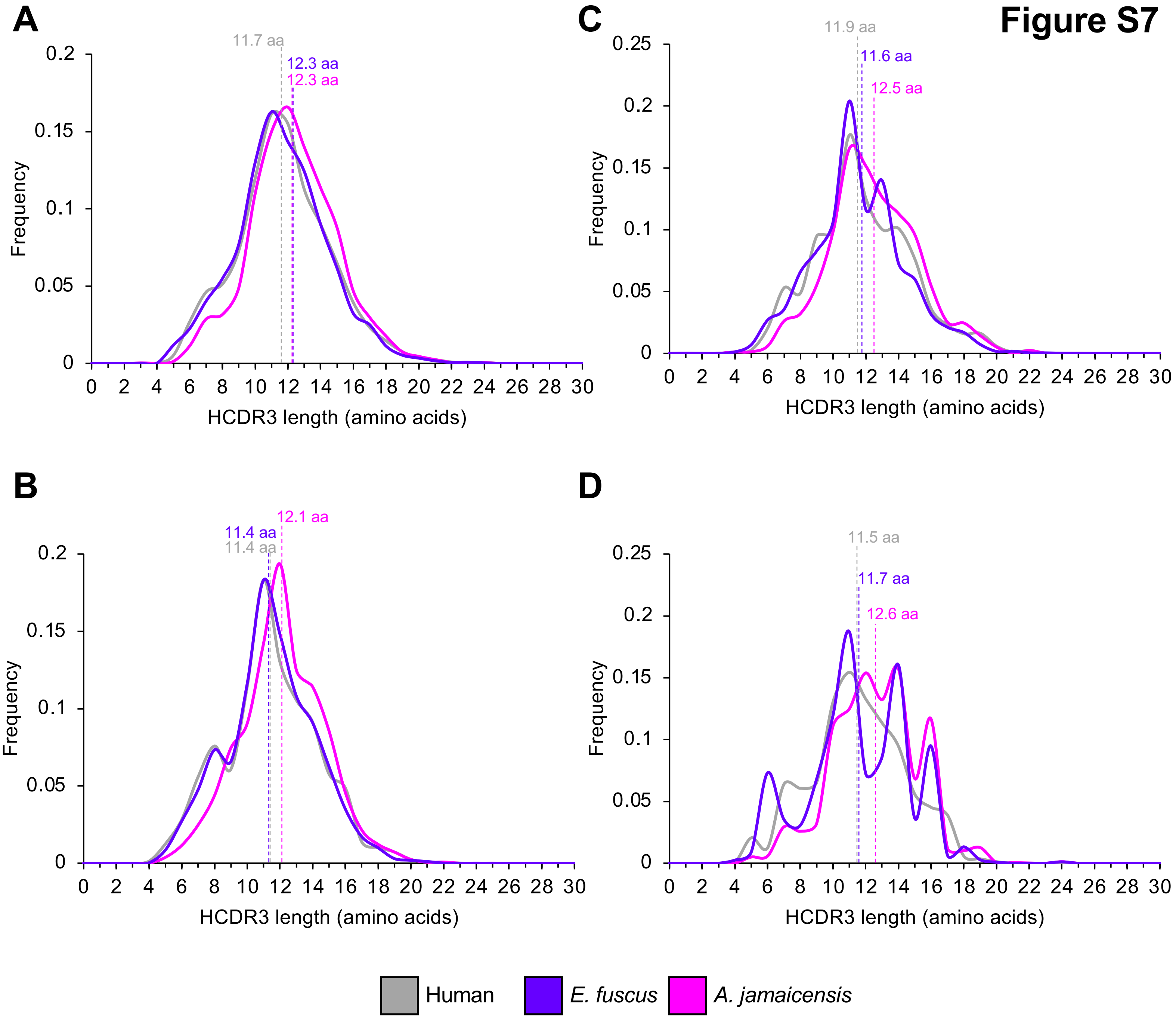

A
11.7 aa
12.3 aa
12.3 aa
Frequency
HCDR3 length (amino acids)
B
11.4 aa
11.4 aa
12.1 aa
Frequency
HCDR3 length (amino acids)
C
11.9 aa
11.6 aa
12.5 aa
Frequency
HCDR3 length (amino acids)
D
11.5 aa
11.7 aa
12.6 aa
Frequency
HCDR3 length (amino acids)
Human
E. fuscus
A. jamaicensis

### Figure S8

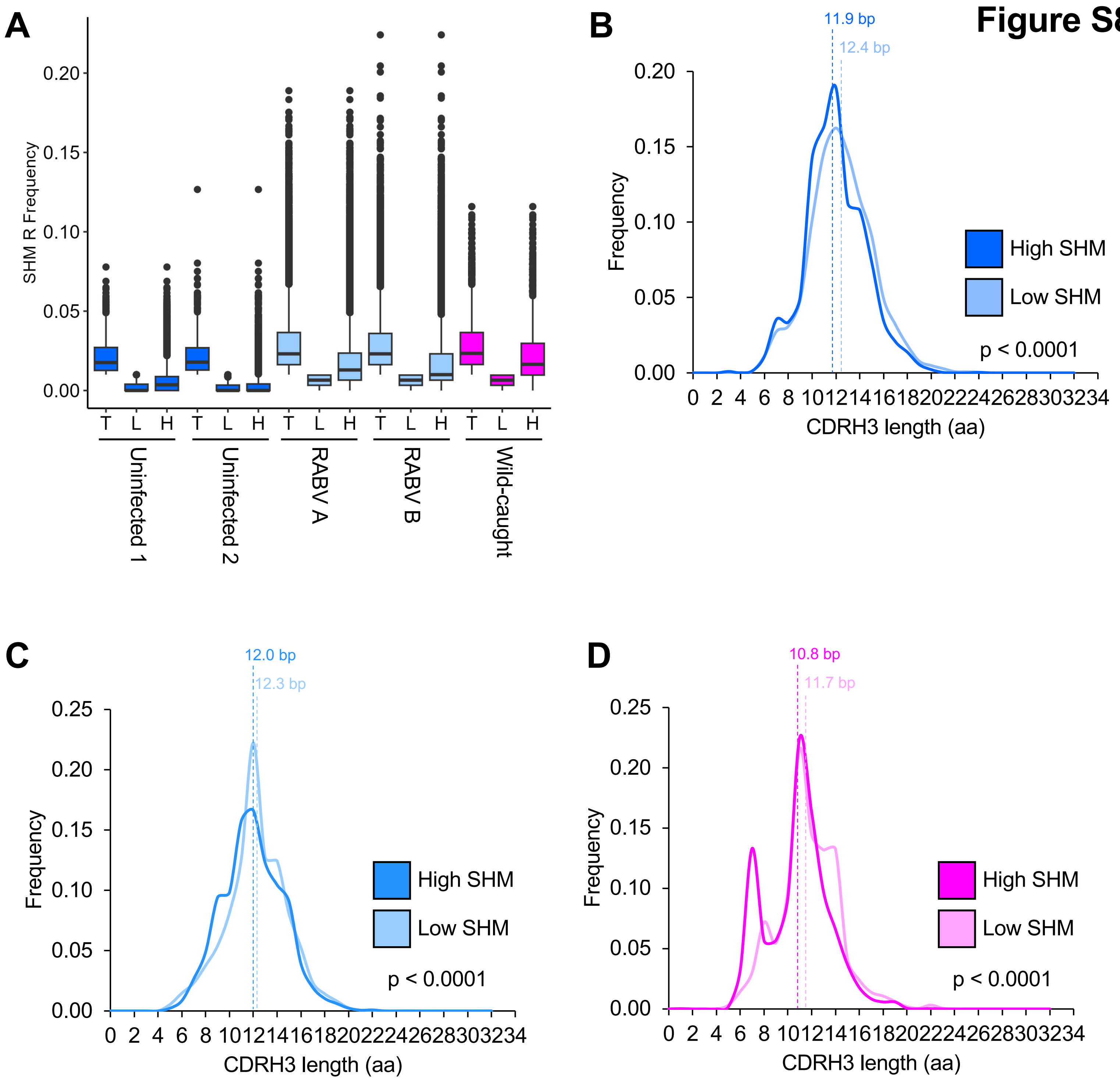
A
SHM R Frequency
0.20
0.15
0.10
0.05
0.00
T L H T L H T L H T L H T L H
Uninfected 1
Uninfected 2
RABV A
RABV B
Wild-caught
B
11.9 bp
12.4 bp
Frequency
0.20
0.15
0.10
0.05
0.00
0 2 4 6 8 10 12 14 16 18 20 22 24 26 28 30 32 34
CDRH3 length (aa)
High SHM
Low SHM
p < 0.0001
C
12.0 bp
12.3 bp
Frequency
0.25
0.20
0.15
0.10
0.05
0.00
0 2 4 6 8 10 12 14 16 18 20 22 24 26 28 30 32 34
CDRH3 length (aa)
High SHM
Low SHM
p < 0.0001
D
10.8 bp
11.7 bp
Frequency
0.25
0.20
0.15
0.10
0.05
0.00
0 2 4 6 8 10 12 14 16 18 20 22 24 26 28 30 32 34
CDRH3 length (aa)
High SHM
Low SHM
p < 0.0001

**Figure S9**

**A**

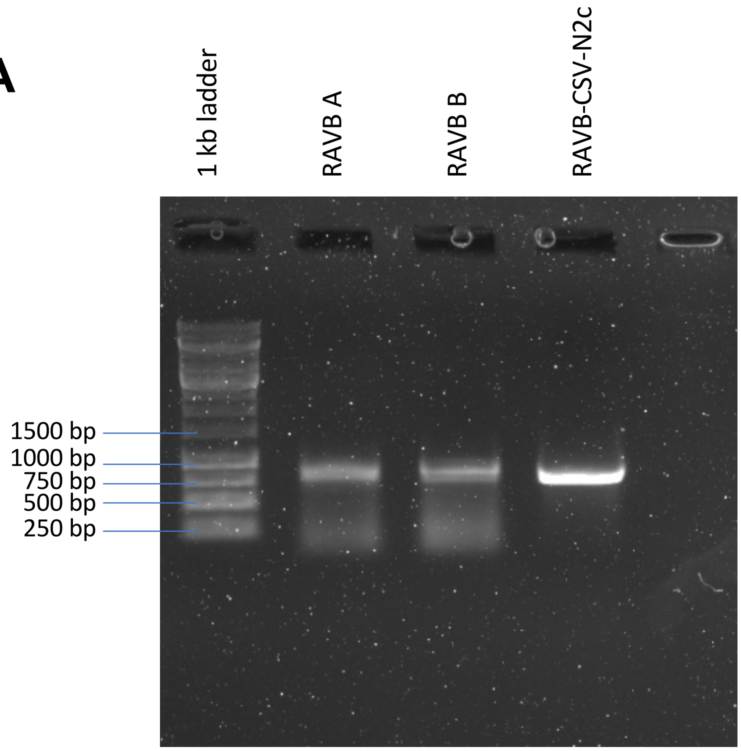
1 kb ladder
RAVB A
RAVB B
RAVB-CSV-N2c
1500 bp
1000 bp
750 bp
500 bp
250 bp

**B**

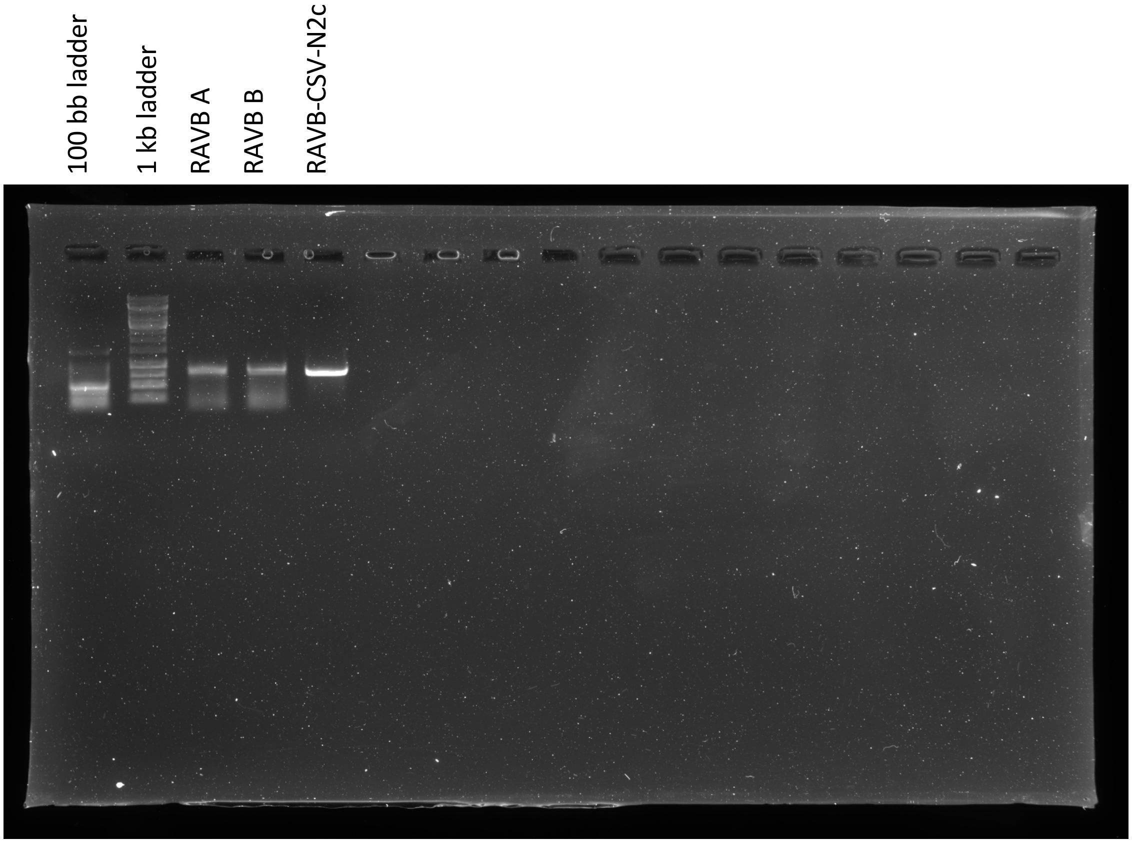
100 bb ladder
1 kb ladder
RAVB A
RAVB B
RAVB-CSV-N2c
